# Mechanisms of MCM2-7 helicase activation and initial DNA melting at near base-pair resolution

**DOI:** 10.64898/2026.01.27.699880

**Authors:** Christopher Weekes, Lia Willerding, Sanjay P. Khadayate, Korbinian Liebl, Audrey Mossler, Alex Montoya, Vanessa Rauthe, Mohammad M. Karimi, Martin Zacharias, Helle D. Ulrich, Christian Speck, L. Maximilian Reuter

## Abstract

During eukaryotic DNA replication initiation, inactive MCM2–7 double-hexamers assembled at replication origins must be converted into two active CMG helicases, yet how this transition is coupled to origin DNA unwinding in vivo remains unclear. Here, we identify a DNA-bound intermediate with an extended genomic footprint that forms during helicase activation. Genome-wide mapping of initial strand separation reveals that DNA unwinding initiates near the N-terminal interface of opposing MCM2–7 hexamers. At these sites, the origin DNA exhibits a conserved AT-rich/GC-rich/AT-rich sequence architecture on which the helicase complex is centred, consistent with a role in promoting DNA extrusion following double-hexamer separation. We further show that restricting hexamer rotation and splitting delays release of the Cdc45-loading factor Sld3, demonstrating that mechanical transitions during helicase activation are tightly coupled to complex disassembly. Finally, we provide in vivo evidence that single-stranded DNA is ejected through a specialised DNA exit gate at the Mcm2/5 interface during helicase activation, which is dispensable for ongoing DNA synthesis. Together, these findings establish a mechanistic framework for how replication origins are remodelled to initiate DNA replication and reveal key intermediates and DNA transactions during helicase activation.

## INTRODUCTION

Replication origins serve as central sites for origin recognition, helicase loading, initial DNA unwinding and firing of origins^1–3^. The accurate regulation of these events is central to genome stability^4,5^. During early G1-phase, ORC, with the help of Cdc6 and Cdt1, loads two copies of the hetero-hexameric MCM2-7 helicase onto double-stranded DNA (dsDNA), a process termed origin licensing. Before it is loaded onto DNA, MCM2-7 adopts a spiral shape with an opening at the Mcm2/5 interface^6,7^, which is used for DNA insertion, and subsequently becomes fully closed in an Mcm4 ATP-hydrolysis dependent fashion^8^. Once the MCM2-7 double-hexamer (DH) forms on DNA, it is initially inactive for DNA unwinding (Supplementary Fig. 1a)^9^. During the G1-S-phase transition, Sld3, Sld7, and Cdc45 bind to the MCM2-7 DH in a DDK kinase-dependent manner (Supplementary Fig. 1b), followed by CDK kinase-dependent binding of Sld2-Dpb11-GINS-Polε (Supplementary Fig. 1c)^10^. This results in the formation of two Cdc45-MCM2-7-GINS-Polε (CMGE) complexes, representing the replisome’s core (Supplementary Fig. 1d)^11^. Mcm10 recruitment then promotes helicase activation (Supplementary Fig. 1e)^12^. These reactions transform the MCM2-7 DH into two individual CMGE hexamers, each encircling single-stranded DNA (ssDNA), generating the basis for bidirectional replication forks^1–3,13^. Consequently, the two CMGEs pass each other, moving head-to-head (Supplementary Fig. 1f), followed by Polδ-mediated initiation of DNA synthesis on the leading strand^14^.

Replication origins are the sites of MCM2-7 loading^10^. They contain specific elements termed A, B1, and B2^15^. The A and B1 elements are binding sites for ORC during the loading of the first MCM2-7 hexamer, while the B2 element serves as the binding site of the second MCM2-7 hexamer^16–20^. Once the MCM2-7 DH is loaded onto the replication origin, it sits on the B1 element and covers large parts of the A and B2 elements^19,20^. Whether DNA unwinding occurs within one of these elements or outside of the origin is not known (Fig. 1a(i and ii)). Moreover, it is unknown whether MCM2-7 hexamers separate by a greater distance during the helicase activation process (Fig. 1a(iii and iv)), which seems relevant as they can be joined across large distances during helicase loading^19,21^, or they remain proximal to their loading site on the B1 element.

**Fig. 1:**
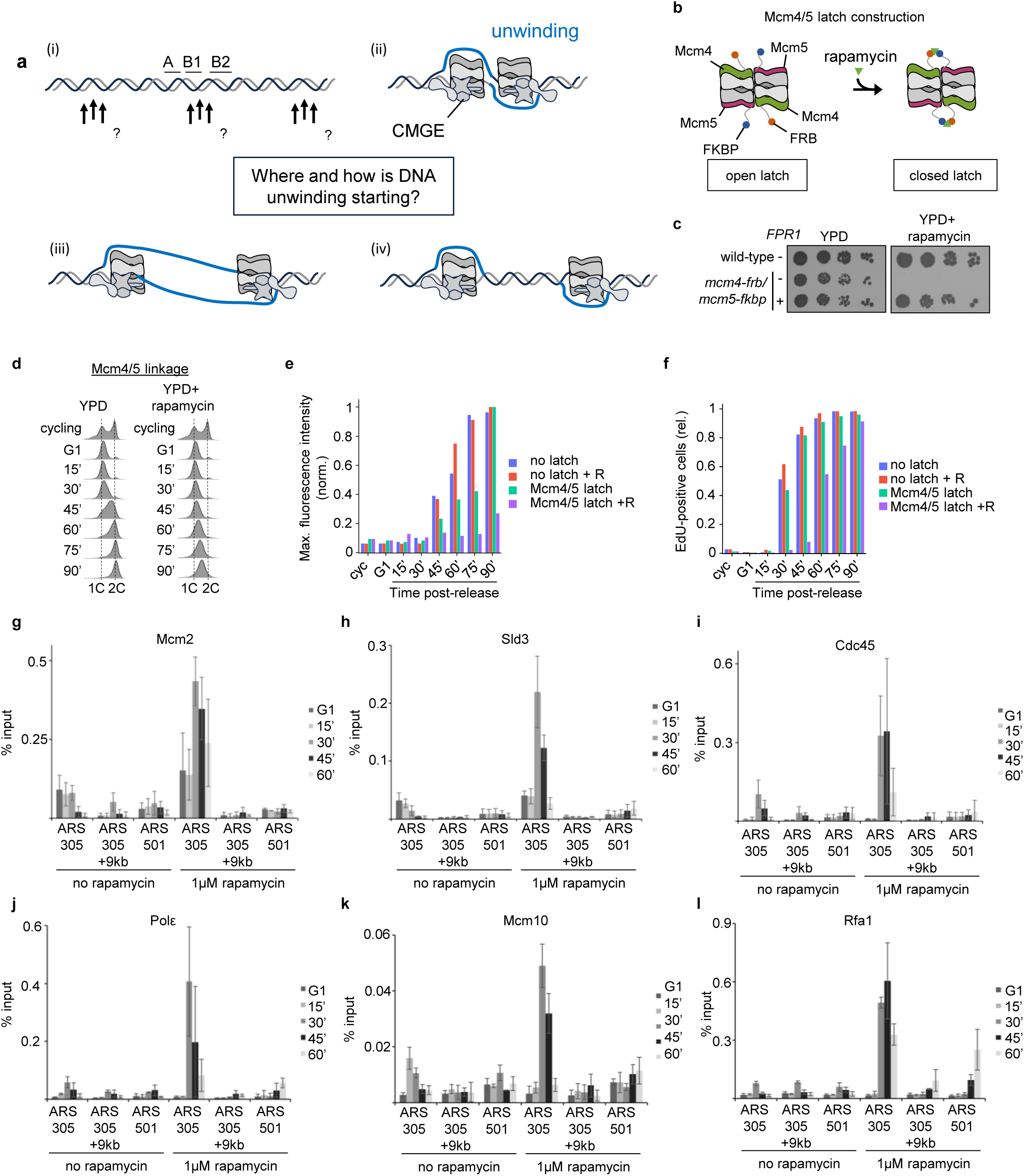
An Mcm4/5 N-terminal latch impairs the release of Sld3 and delays helicase splitting. (**a**) The location of initial DNA unwinding on origin DNA (i) and potential models of DNA unwinding (ii-iv). (**b**) Concept of the latch construct. Integration of FRB and FKBP domains with flexible linkers into the N-termini of MCM4 and MCM5 was used to produce an inducible, inter-hexameric linkage *in vivo* that can be conditionally closed by rapamycin. (**c**) Dot spot assay of the Mcm4/5 linkage mutant on full media (YPD) or YPD-rapamycin (100 nM) in the presence or absence of an extrachromosomal copy of *FPR1.* (**d**) Representative flow cytometry profiles of a strain with a Mcm4/5 linkage in the absence (left panel) and presence of rapamycin (1 µM, right panel). Cycling cells were arrested for 2 hrs and treated with rapamycin before release into S-phase for 90 minutes. (**e** and **f**) EdU incorporation analysis to visualise (**e**) normalised EdU signal intensity and (**f**) the percentage of cells positive for EdU incorporation. α-factor arrested cells were released in S-phase in media containing 2 µM EdU in the absence or presence of rapamycin (+R, 1 µM) for the indicated times. Cycling cells without EdU treatment served as background control. (**g**-**i**) ChIP-qPCR profiles of various strains harbouring the Mcm4/5 linkage. Evaluation of (**g**) Mcm2, (**h**) Sld3, (**i**) Cdc45, (**j**) Polε, (**k**) Mcm10, and (**l**) Rfa1 recruitment to *ARS305*, *ARS305*+9kb and *ARS501*. Mcm4/5 linkage strains were treated with or without rapamycin (1µM) and harvested at the indicated timepoints after α-factor arrest and release at 25 °C for the indicated amounts of time. Presented are the average and standard deviation of at least three biological replicates.

Recent structural insights have revealed that, upon CDK-dependent helicase activation, the two hexamers partially separate, accompanied by a 100° rotation of the two hexamers and a minor destabilisation of the DNA, but remain closely associated (Supplementary Fig. 1d)^22^. Then, dual Mcm10 interactions with the MCM2-7 complex, involving the N-terminal MCM interface, separate the two hexamers^23,24^. This is associated with DNA strand extrusion starting from the N-terminal region of MCM2-7, followed by complete DNA unwinding as the helicase pore narrows^24^. Moreover, after the initial DNA unwinding, the two hexamers pass each other^25^, and it has been suggested that pulling of DNA by both hexamer motors facilitates lagging-strand ejection/extrusion^26^. Once the two hexamers have passed each other, the N-terminal face of these CMGs becomes freely accessible, thereby allowing binding of the fork protection complex, consisting of Tof1, Csm3, and Mrc1^27^.

However, many steps in helicase activation remain incompletely understood. Is the rotation of the two hexamers and their separation coordinated with complex assembly and the release of activation factors? Do all six potential MCM2-7 interfaces function similarly for initial DNA unwinding and ongoing DNA synthesis, as observed during DPC bypass^28^, or is one more important? Is it possible to detect the footprint of the CMG helicase when it becomes activated? Where does the initial DNA unwinding occur within the replication origin? Are specific DNA features associated with DNA unwinding, and where does the unwound DNA appear relative to the two MCM2-7 complexes (Fig. 1a)?

The overall technical framework of this study is to analyse helicase activation *in vivo* using a combination of chemical genetics, genomics, and proteomics. By interfering with inter-hexamer rotation, DH separation, and DNA extrusion via rapamycin-dependent linkages between neighbouring Mcm subunits, we were able to study these events *in vivo*.

Interestingly, we discovered that initial DNA strand separation during helicase activation occurs near the ORC-DNA-binding site, is characterised by a conserved AT-rich/GC-rich/AT-rich architecture, arises near/in between the interfaces of the two MCM2-7 hexamers, is associated with Sld3 release and utilises a specific MCM2-7 DNA exit gate. As such, our work highlights how replication factors function and utilise replication origins to activate genome-wide helicase activity, providing a comprehensive *in vivo* framework for understanding the regulation of these processes and their contribution to genome stability.

## RESULTS

### Separation and rotation of the two MCM2-7 hexamers are linked to the Sld3 release

Helicase activation is a multi-step process. Cryo-EM analysis has linked CDK-dependent CMG assembly to the initial separation of DHs and to MCM2-7 rotation^22^. However, how inhibition of DH separation or MCM2-7 rotation affects complex assembly and disassembly *in vivo*, is unknown. Here, we have employed a rapamycin-dependent linkage system between two MCM2-7 hexamers to interrogate how inhibition of MCM2-7 inter-hexamer rotation and separation of the two hexamers affects the process (Fig. 1b).

Within the MCM2-7 DH, Mcm4 from one hexamer is positioned proximal to Mcm5 from the other hexamer, and this pairing is repeated for the subunits across the other side of the complex. By integrating an FRB domain and linkers in the N-terminus of Mcm4, and an FKBP domain and linkers in the N-terminal domain of Mcm5, two separate “latches” were generated on either side of the DH (Fig. 1b). Upon addition of rapamycin, dimerisation of these domains would create links between the two hexamers and close the latch, thereby affecting inter-hexamer rotation and DH separation. Indeed, the rapamycin-induced formation of the McmN4/5 latch led to a growth arrest in the strain (Fig. 1c). Re-introducing the previously deleted *FPR1* gene coding for an endogenous yeast rapamycin-binding protein in the latch strain rescued cell growth under rapamycin treatment (Fig. 1c), highlighting that it generates an Mcm2-linked FRB-*FPR1* interaction that interferes with the FKBP-FRB interaction. Thus, the Mcm2-FFRB-*FPR1* connection does not affect cell growth, but the FKBP-FRB and rapamycin-mediated closing of the Mcm4-Mcm5 latch induces a growth arrest. To assess the impact of the closed latch on the cell cycle, we induced a G1 arrest with α-factor and then released the cells synchronously into the cell cycle in the presence and absence of rapamycin. While untreated cells progressed normally into S-phase, rapamycin treatment induced a cell-cycle slowdown (Fig. 1d). Here, we observed a concentration-dependent cell-cycle slowdown, with lower concentrations producing weaker phenotypes presumably due to latch breaking (Supplementary Fig. 2a and b). Furthermore, we observed a rapamycin-dependent block in EdU incorporation up to 60 minutes post α-factor release, which was followed by almost all cells incorporating a low level of EdU at the 90-minute time point (Fig. 1d-f, Supplementary Fig. 2c). However, even after 90 minutes, cells had only replicated ∼30% of their DNA (Fig. 1e). As such, the latch connecting Mcm4 with Mcm5 strongly hinders cell cycle progression from the G1-phase into the S-phase up to 60 minutes, followed by very slow progression through S-phase. We interpret this as the linkage breaking at a subset of origins beginning at approximately 60 minutes, which could explain the subsequent increase in DNA synthesis.

To understand which step in helicase activation is affected, we measured complex assembly by ChIP-qPCR, as previously used to study the function of Mcm10^29–31^. We focused our analysis on the early time points, since our time-course data (Fig. 1f) suggested a breakage of the latches at later time points, e.g. 60 minutes onwards, as an increase in EdU-positive cells can be detected (Fig. 1f). Here, we observed specific recruitment of MCM2-7 to replication origins. In the presence of rapamycin, DNA binding at ARS305 was strongest at 30 minutes (Fig. 1g). This was reminiscent of the increased ChIP efficiency previously observed in the context of helicase activation in the absence of Mcm10^32^. We suggest two reasons for the increased ChIP signal: 1) the latch-mediated arrest renders the CMG complex stationary at the origin, while in the absence of rapamycin, the forks leave the origin quickly, diminishing the ChIP signal at the origin (Fig. 1g, compare ARS305+9kb signals in untreated and rapamycin-treated conditions). 2.) CMG-mediated DNA unwinding generates ssDNA-protein contacts, which react more easily with formaldehyde than dsDNA-protein contacts. From the 45-minute time point onwards, the MCM2-7 (Mcm2) signal decreased (Fig. 1g), which may indicate reorganisation of the complex or partial disruption of the latches. Interestingly, Sld3 associates early with MCM2-7 but becomes stabilised in the presence of rapamycin at the 30-minute timepoint and remains partially associated even when most Sld3 in the rapamycin-lacking reaction has already been released (Fig. 1h). Thus, our data show that Sld3 is stabilised by the closed Mcm4/5 latch. Cdc45 recruitment is observed from the 30-minute time point onward (Fig. 1i). In the context of rapamycin, we observed a stabilisation of Cdc45 similar to that of Mcm2 (compare Fig. 1i with 1g), and Polε showed similar recruitment (Fig. 1j). Finally, in the presence of rapamycin, we observed delayed and increased binding of Mcm10 to replication origins (Fig. 1k) and a major rapamycin-dependent delay and stabilisation of RPA (Fig. 1l). The data are consistent with CMG formation and initial DNA unwinding. However, the latch construct resulted in increased Sld3 binding and delayed Mcm10 binding. We thus conclude that the closed latch interferes with the replication fork assembly process at a late activation stage. Therefore, our results indicate that hexamer rotation and complete DH splitting are necessary for efficient Sld3 release and timely Mcm10 binding.

### The Mcm2/5 linkage causes an arrest in replication fork assembly

During helicase loading, dsDNA is inserted through the partially open Mcm2-Mcm5 interface into the MCM2-7 ring^6,7,33^. Loading of two MCM2-7 hexamers generates an MCM2-7 DH encircling dsDNA, which is inactive for DNA unwinding^34,35^. In S-phase, Mcm10 promotes the extrusion of ssDNA from the helicase core^24,30,31,36^. However, the underlying mechanisms of MCM2-7 ring opening during S-phase remain elusive. In particular, it needs to be clarified whether a specific Mcm interface is essential for helicase activation, or whether all six interfaces can contribute equally to strand extrusion as observed during DPC bypass *in vitro* (Fig. 2a)^28^. Therefore, we set out to systematically characterise all six Mcm interfaces for their ability to capture extruded ssDNA from the MCM2-7 ring during helicase activation *in vivo*, using rapamycin-inducible FRB/FKBP linkages between MCM subunits (Fig. 2b)^33^. We endogenously integrated the FRB/FKBP domains at all neighbouring MCM2-7 interfaces, targeting flexible loops on the protein surface. To obtain a more holistic insight, we generated rapamycin-inducible FRB/FKBP linkage at both the N-terminal and C-terminal domains of neighbouring Mcm subunits (Fig. 2c and d). For interfaces involved in DNA insertion/extrusion, rapamycin-dependent lethality should be observed at both N- and C-terminal positions upon linkage. In contrast, lethality observed only upon linkage at either the N- or C-terminal domains would indicate an essential function distinct from DNA insertion/extrusion for the specific interface. We found that the linkage, which incorporated FRB and FKBP moieties in the N-terminal sections of Mcm2/5 interface and the Mcm5/3 interface produced a lethal phenotype in the presence of rapamycin (Fig. 2c)^33^. However, FRB and FKBP moieties only produced a lethal phenotype at the C-terminal sections of the Mcm2/5 interface in the presence of rapamycin (Fig. 2d). Thus, the data are consistent with the Mcm2/5 interface functioning in DNA insertion and DNA extrusion, while the data for the Mcm5/3 interface point towards a different role. Indeed, a diversion tunnel, feeding the lagging strand to Polδ, was identified between the zinc fingers of Mcm3 and Mcm5 in the unwinding CMG^37^.

**Fig. 2:**
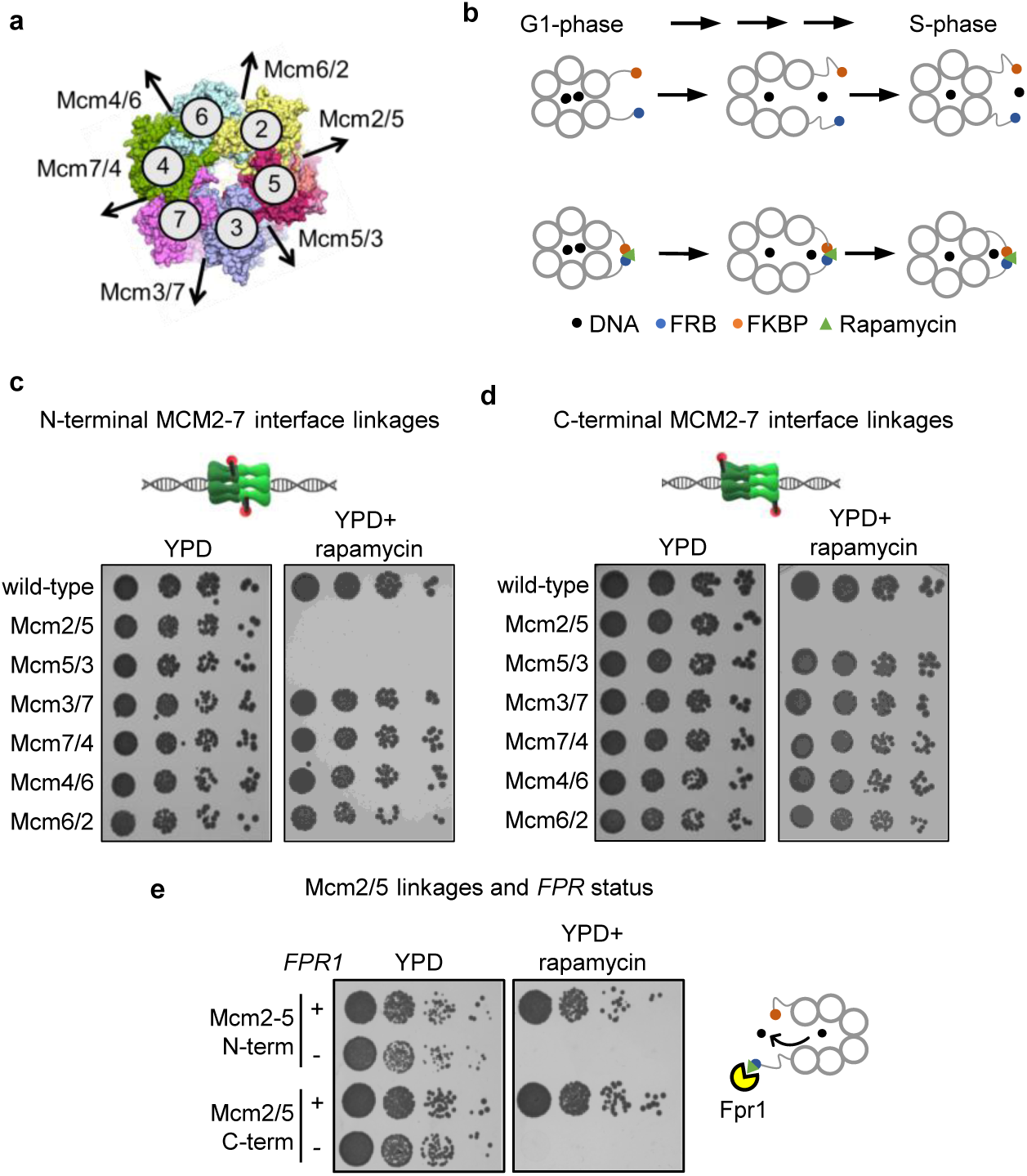
Construction and evaluation of MCM2-7 intra-hexameric linkages. (**a**) Rationale of the MCM2-7 intra-hexameric linkage construction. The helicase comprises six unique interfaces, of which any could be used for ssDNA extrusion during helicase activation. (**b**) Schematic logic of a linkage construct integrated into the natural MCM loci. FRB/FKBP domains with flexible linkers (blue and red dots with lines) were inserted in flexible regions of all Mcm subunits that produce an intra-hexameric interface within the MCM2-7 helicase. In the absence of rapamycin, ssDNA will be released normally. In contrast, when a linkage is established through the addition of rapamycin in G1-phase (green), extruded ssDNA will be captured by rapamycin-induced linkage during S-phase onset. This approach can unveil which and how many interfaces can be utilised to extrude ssDNA during helicase activation. (**c** and **d**) All (**c**) N-terminal and (**d**) C-terminal combinations of linkages between neighbouring Mcm interfaces were constructed and tested for viability in the absence and presence of rapamycin. The Mcm2/5 inducible linkages are uniquely lethal (both N- and C-terminal linkages) within the MCM2-7 complex *in vivo*. Dot spot assay of all linkage mutants on full media (YPD) or YPD-rapamycin (100 nM). (**e**) *FPR1* expression reverses the rapamycin-induced lethality in any Mcm2/5 linkage mutant strain. Dot spot assay of the N-terminal and C-terminal Mcm2/5 linkage mutant strains on full media (YPD) or YPD-rapamycin (100 nM) in the presence or absence of an extrachromosomal copy of *FPR1*.

Re-introducing the *FPR1* gene coding for an endogenous yeast rapamycin-binding protein, in the N- and C-terminal Mcm2/5 linkage-containing strains, rescued cell growth under rapamycin treatment (Fig. 2e; Supplementary Fig. 3a). These results demonstrate that neither the FRB/FKBP domains by themselves nor rapamycin itself affects cell growth in the context of the Mcm2/5 linkage strains.

As it is already known that the Mcm2/5 interface functions in helicase loading^33^, we then focused on the role of the Mcm2/5 interfaces during the G1/S transition. To address this question, we arrested cells in G1-phase, added rapamycin to form the FRB/FKBP linkage and released cells synchronously into the cell cycle (Fig. 3a). Indeed, the rapamycin-induced linkage at the Mcm2/5 interface at the N-terminal and C-terminal domain hindered the G1/S transition while untreated cells progressed through the cell cycle (Fig. 3b and c). However, treatment of the N- and C-terminal Mcm2/5 linkage strain after the release into S-phase had only a minor impact on bulk DNA synthesis (Supplementary Fig. 3b and c). Thus, we conclude that the Mcm2/5 linkage primarily affects helicase activation *in vivo*.

**Fig. 3:**
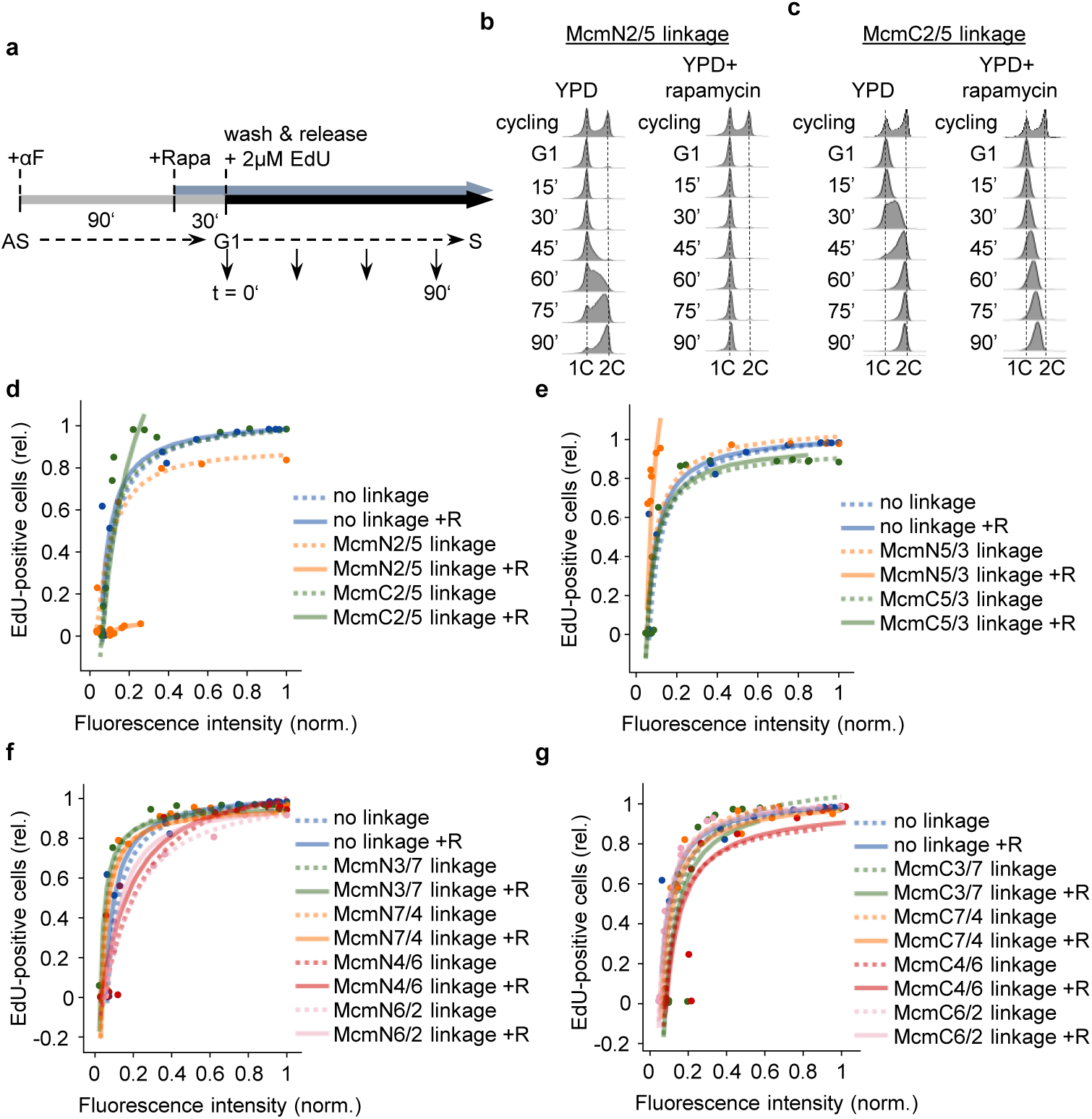
The Mcm2/5 linkages arrest helicase activation at different stages in the G1-S-phase transition. (**a**) Experimental scheme for the EdU incorporation time-course experiments. Cells arrested in G1 phase with α-factor (αF) were treated with or without rapamycin (Rapa) before release into the cell cycle in the presence of 2 µM EdU. Samples were taken in 15-minute intervals for 90 minutes at 25 °C. (**b** and **c**) Representative flow cytometry profiles of a strain with an (**b**) N-terminal or (**c**) C-terminal Mcm2/5 linkage in the absence and presence of rapamycin (100 nM). (**d**-**g**) EdU incorporation time course experiments reveal cell-cycle progression differences in the Mcm2/5 linkage mutant strains. Composite plots of EdU incorporation efficiency measures (EdU positive cells vs. normalised EdU signal intensity) of strains harbouring (**d**) an McmN2/5 linkage, (**e**) an McmC2/5 linkage, (**f**) all other N-terminal, and (**g**) C-terminal Mcm linkages in the absence (dotted lines) and presence of rapamycin (+R, 100 nM, solid lines). Strains are colour-coded, and individual dots represent the relative percentage of EdU-positive cells and the geometric mean of EdU fluorescence intensity of at least 40,000 cells at a given time point (cyc, G1-90 min).

By arresting cells in G1-phase, releasing them into the cell cycle, and then measuring EdU incorporation and DNA content throughout the cell cycle (Fig. 3a-g), we observed a rapamycin-dependent block in EdU incorporation for the N-terminal Mcm2/5 linkage construct (Fig. 3b and d), suggesting an arrest before the initiation of DNA synthesis. The C-terminal linkage, on the other hand, resulted in a near-normal increase of EdU-positive cells, but these cells failed to continuously synthesise DNA (Fig. 3c and d), suggesting normal initiation and a block in elongation. The N-terminal Mcm5/3 linkage construct also resulted in a near-normal increase of EdU-positive cells, but these cells failed to synthesise DNA as well (Fig. 3e and Supplementary Fig. 3d). In contrast, all other linkage constructs (including the C-terminal Mcm5/3 linkage) displayed normal EdU incorporation and DNA incorporation in the presence of rapamycin (Fig. 3f and 3g, Supplementary Fig. 4). Based on that analysis, the Mcm2/5 C-term linkage is consistent with normal strand separation and a block in elongation. In contrast, the N-terminal Mcm2/5 linkage appears to arrest helicase activation at an earlier stage. The N-terminal Mcm5/3 construct initiated DNA replication, but this was more limited in comparison to the Mcm2/5 C-terminal construct. Based on these data, we continued our work with the Mcm2/5 C-terminal construct, which indicated near-normal helicase activation and a defect in DNA synthesis.

### Assembly of a late helicase activation intermediate is linked to DNA extrusion

To understand how replisome complex assembly is affected by the Mcm2/5 linkage, we employed chromatin immunoprecipitation coupled to mass spectrometry (ChIP-MS) and analysed the C-terminal Mcm2/5 linkage construct and a corresponding control strain identical to it except for the absence of the Flag-tag on Mcm4.

In G1-phase, we found strong enrichment of MCM2-7 proteins in the Mcm2/5 linkage strain in comparison to the control strain, while recruitment of helicase activation and DNA synthesis factors was reduced, as expected (Fig. 4a). At the 30-minute post-release time point, enrichment of helicase-activation factors was observed (Fig. 4b). By comparing the Mcm2/5 linkage strain 30 minutes post-G1 release with the G1-phase arrested strain, we found particularly strong recruitment of Cdc45, Sld2, Dpb11, and Mcm10 at the 30-minute time point (Fig. 4c). Furthermore, when comparing the Mcm2/5 linkage strain with the strain missing the linkage 30 minutes post-G1 release, we detected strong enrichment of Sld2, Dpb11, and Mcm10, while Tof1 was the most de-enriched factor (Fig. 4d).

**Fig. 4:**
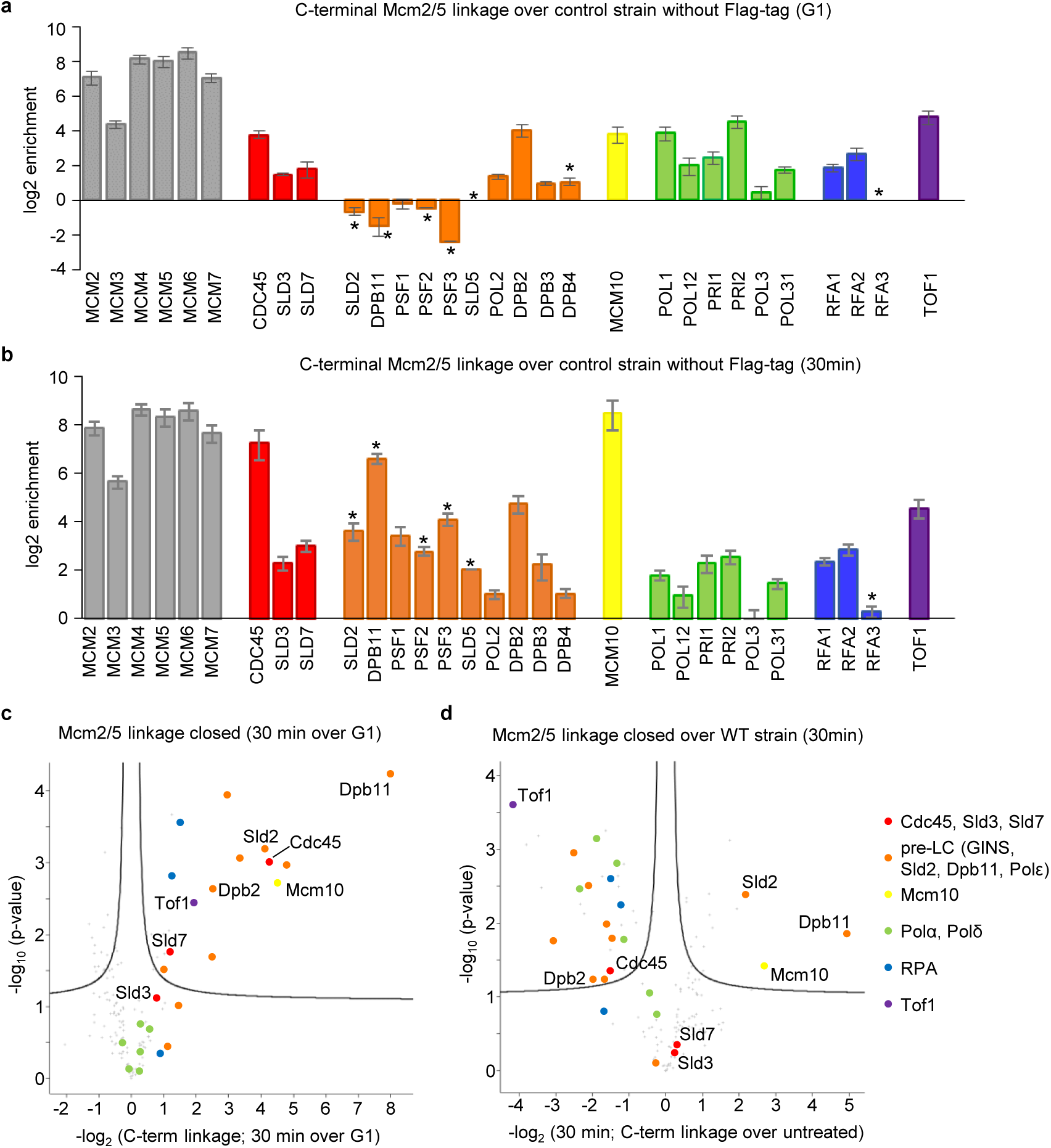
McmC2/5 linkage-containing complexes arrest at a late helicase activation stage. MCM2-7 complexes arrested by the McmC2/5 linkage enrich for replication activation and fork-associated proteins. (**a**) Mcm4 ChIP-MS enrichment of core replisome factors in the Mcm2/5 C-terminal linkage strain in G1-phase. (**b**) Mcm4 ChIP-MS enrichment of core replisome factors in the Mcm2/5 C-terminal linkage strain released into S-phase for 30 min. Enrichment values are given as the log_2_ of the mean LFQ enrichment of subject proteins in the Mcm4-5xFLAG strain (IP) over a negative control (NC) strain from four independent replicates. Asterisks mark proteins that lacked intensity measurements in the NC. Error bars represent the standard error (S.E.) of the non-logarithmic mean. (**c** and **d**) The Mcm2/5 C-terminal linkage construct supports assembly of a late helicase activation intermediate. (**c**) Volcano plot summarising replication factor enrichments of Mcm4-containing complexes from cells released into S-phase for 30 min versus G1-arrested cells in the context of the closed Mcm2/5 linkage. Important replication initiation factors are colour-coded: Cdc45, Sld3/7 (red), pre-LC components (orange), Mcm10 (yellow), Pol1 and 3 (green), RPA (blue) and Tof1 (purple). (**d**) The McmC2/5 gate linkage arrests MCM2-7 helicase activation at a late stage. Volcano plot summarising replication factor enrichments of Mcm4-containing complexes from closed McmC2/5 linkage cells over wild-type cells released into S-phase for 30 min. Replication initiation factors are colour-coded as in (c).

Mcm10 is involved in ssDNA extrusion and in splitting the MCM2-7 DH^23,24^. It separates the two hexamers held together at their N-terminal ends into two complexes, each encircling one ssDNA strand^10,24^. Once the two hexamers have passed each other, Tof1 can interact with the N-terminal face of MCM2-7^27^. Significantly, we observed enrichment of Mcm10 and the absence of Tof1, suggesting arrest after Mcm10 association but before Tof1 binding (Fig. 4d).

These results were further corroborated by assessing the temporal recruitment of proteins to replication origins via ChIP-qPCR (Supplementary Fig. 5). When treated with rapamycin, we observed persistent recruitment of Mcm4, Cdc45, Polε, and Mcm10 at the *ARS305* (early origin) and *ARS501* (late origin) origin consensus sequences. The very strong enrichment at the origin demonstrates that CMG helicase is immobile in the context of the closed Mcm2/5 linkage, although all helicase activation factors are present. Thus, the rapamycin-induced Mcm2/5 linkage enables the assembly of a late immobile replisome intermediate enriched for Sld2, Dpb11, and Mcm10, and depleted of Tof1. Therefore, we conclude that the CMG complex forms, but it cannot mature into a full replisome (Fig. 4d, Supplementary Fig. 6a and b).

### Helicase activation generates a large DNA footprint across the replication origin

Next, we attempted to identify the DNA footprint of the replisome assembly intermediate. Currently, it is unknown how much dsDNA is covered during helicase activation (Supplementary Fig. 1); therefore, it is unclear how much accessible space is required at replication origins for this process. Moreover, we do not know whether a specific origin element is involved, or whether complex assembly occurs outside of the defined origin sequence (Fig. 1a). Initially, we subjected the Mcm2/5 C-terminal linkage strain to ChIP-Exo 5.0 in G1-phase in the presence of rapamycin and identified a 64 bp footprint (Fig. 5a) centred across the B1 element, identical to the wild-type footprint location and AT content distribution that has been previously observed (Fig. 5b, Supplementary Fig. 7a)^20^. 30 minutes post G1 release, we observed in addition two larger footprints (Fig. 5c, Supplementary Fig. 7b). The prominent 72 bp footprint could represent an intermediate, where the two MCM2-7 hexamers are partially separated due to ATP-hydrolysis, resulting in initial strand separation within the CMG, as observed by cryo-EM and in biochemical experiments^25^ (Fig. 5c and Supplementary Fig. 1d). The less abundant 138 bp footprint may represent two separate CMGs, where Mcm10 succeeded in splitting the MCM2-7 DH, and Polε contacts the DNA. This results in an extended DNA footprint, with the closed Mcm2/5 linkage blocking further DNA unwinding (Fig. 5c, Supplementary Fig. 7b). We noted that at 30 minutes post G1 release, the closed Mcm2/5 linkage complex stays associated in proximity of the origin (Fig. 5d) and complexes are enriched at early origins consistent with normal activation dynamics (Fig. 5e)^38^. We conclude that the closed Mcm2/5 linkage arrests helicase activation, generating discrete complexes up to 138 bp in size that are immobile, positioned across origin sequences and require >2 times the space of the MCM2-7 DH, exceeding even the average nucleosome-depleted region of 110-130 bp at replication origins^39^. Moreover, DH splitting is not associated with extensive backtracking or sliding of individual CMG complexes; they remain in direct proximity to the MCM2-7 loading site.

**Fig. 5:**
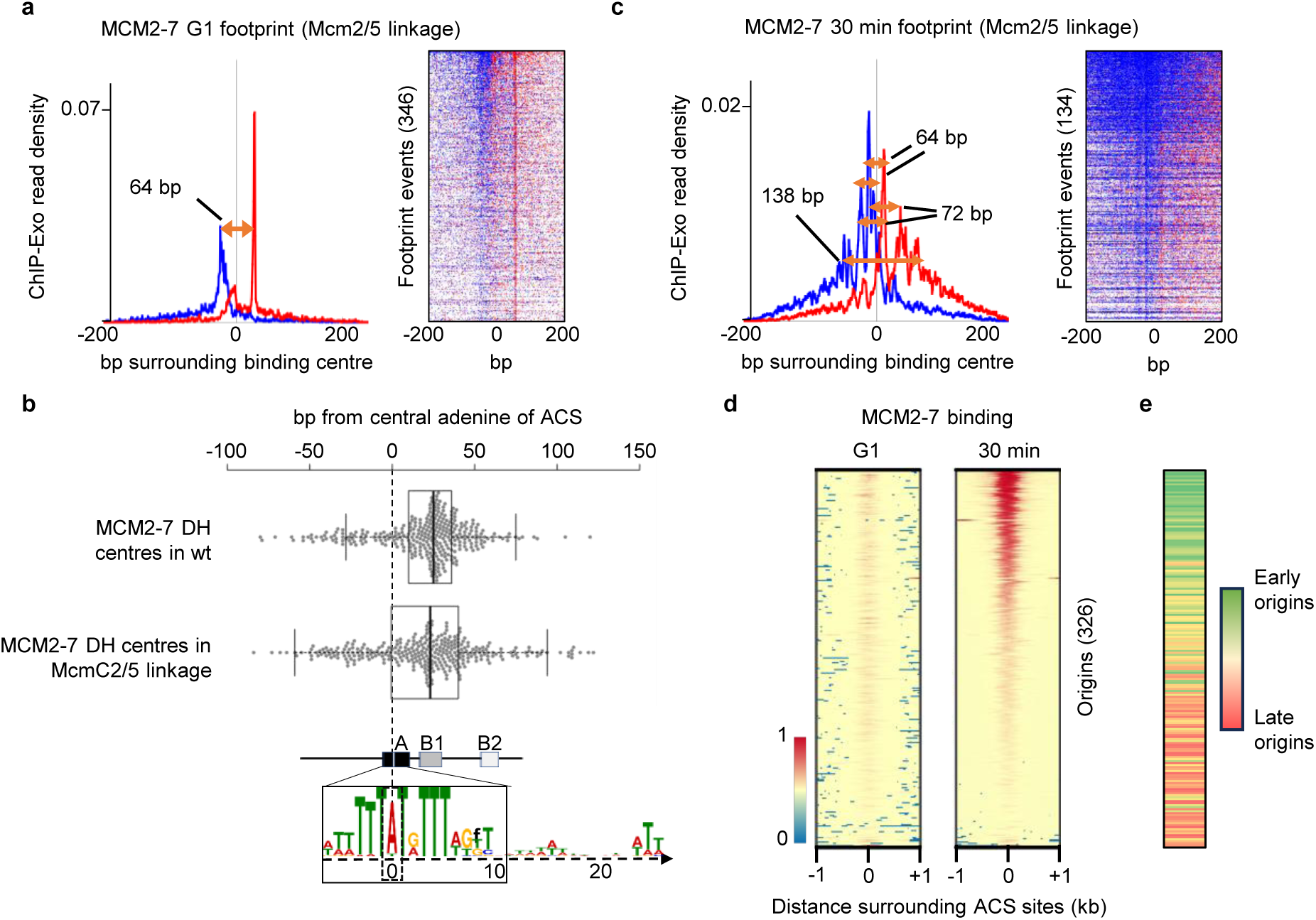
The Mcm2/5 linkage induced footprints during helicase activation. (**a**) Composite plots of ChIP-Exo 5.0 tag distribution patterns (forward strand in blue and reverse strand in red) of Mcm4 from G1-arrested cells. Distinct footprints of MCM2-7 are displayed based on 346/134 confirmed core origins ±200 bp. Arrows indicate footprint sizes (orange arrows). Representative heat maps of ChIP-Exo 5.0 tag enrichments, sorted by decreasing signal, of the MCM2-7 footprints are shown ± 200 bp. (**b**) MCM2-7 DHs from wt and the McmC2/5 linkage strains cover the same area of origins. Distance calculations from the central adenine (dotted line) of the A-element sequences to the binding centre of wt MCM2-7 DHs (n = 329, Reuter et al, 2024 Nature Communications) and MCM2-7 DHs from an McmC2/5 linkage-containing strain (n= 346) are depicted. The locations of conserved origin DNA elements are based on *ARS1* and were aligned with the consensus A-element sequence. Individual distances (dots) as well as the median, 1st, and 3rd quartiles are shown. Whiskers extend to < 1.5x IQR from 1st and 3rd quartiles. (**c**) Composite plots of ChIP-Exo 5.0 tag distribution patterns as in (a) of Mcm4 from cells released for 30 min into S-phase in the context of the closed McmN2/5 linkage. Distinct footprints of MCM2-7 are displayed based on 134 confirmed core origins ±200 bp. (**d**) MCM2-7 complexes expand their footprint when activated. ChIP-Exo 5.0-derived heatmaps of Mcm4 binding to 326 origins at G1 and 30 min post-release in the McmC2/5 linkage strain treated with rapamycin. Centred are A-elements ±1 kb, sorted by Mcm4 signal intensity. (**e**) Expanding MCM2-7 footprints correlate with replication timing. ChIP-Exo 5.0 tag enrichments from (d) were correlated with replication profile data (n= 187; Mueller and Nieduszynski, 2012, Genome Research). Origins are gradient colour-coded from early (green) to late (red) origins.

### DNA sequence properties at sites of replisome assembly

The MCM2-7 DH is the starting point for helicase activation. As the MCM-DNA complex has been structurally characterised, we were able to analyse the conformation of bound origin DNAs within the MCM2-7 DH. We observed a specific twist in the base-pairs near the centre of the protein-DNA complex, indicating that MCM2-7 alters the DNA structure (Fig. 6a). When accounting for the MCM2-7-induced deformation of the DNA, calculations based on an Ising model revealed a localised likelihood for strand-dissociation near the DH interface, regardless of the encircled sequence^40^ (Fig. 6b). This is reminiscent of the human DH complex, which has 3 bp unwound at its centre^41–43^. Thus, the data suggest that the yeast and human MCM2-7 DHs are similar, with DNA-destabilising activity near their centres.

**Fig. 6:**
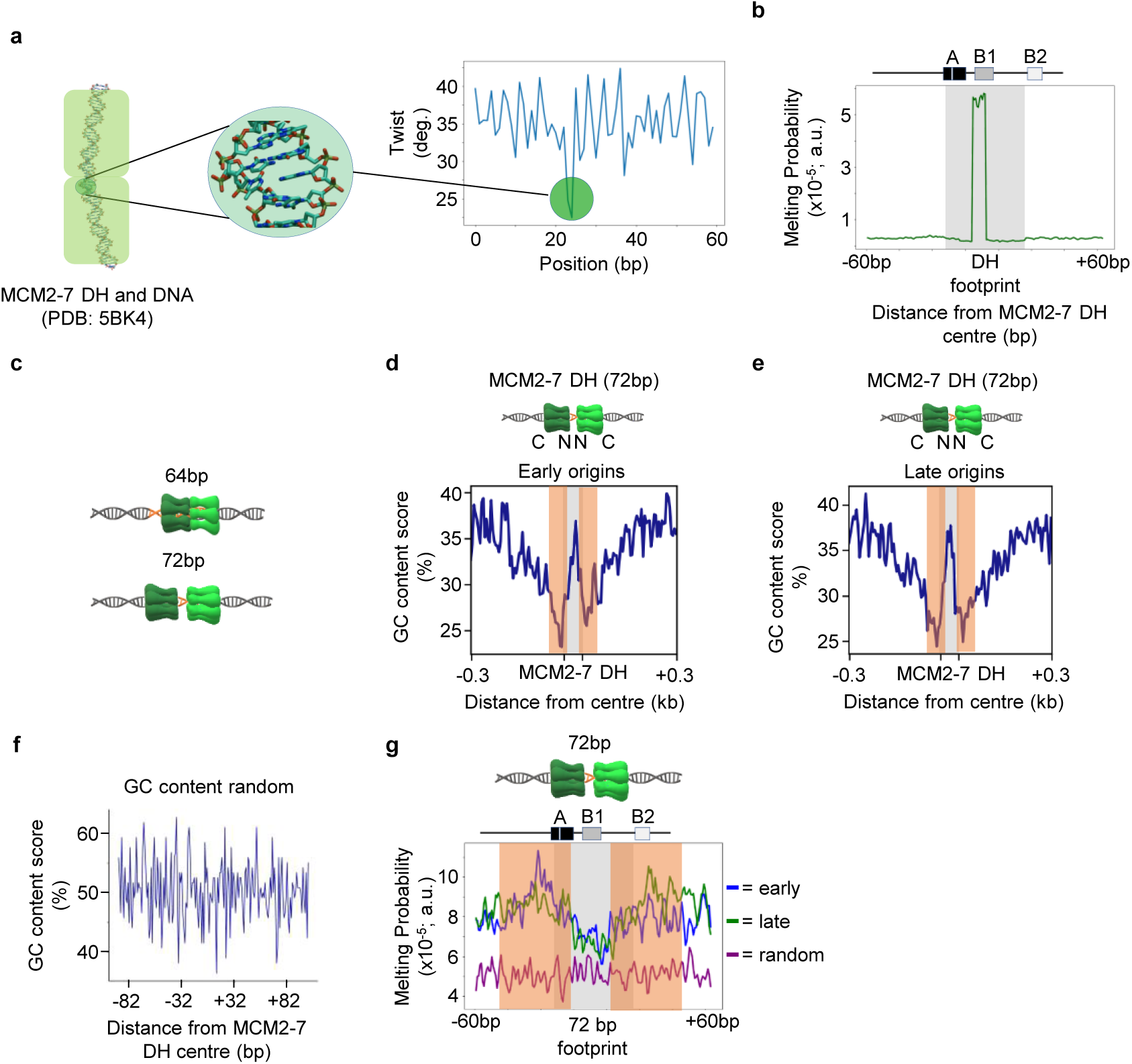
*In silico* analysis of DNA untwisting and GC content in the context of the MCM2-7 DH. (**a**) Analysis of the local untwisting by the MCM2-7 DH encircling dsDNA (PDB ID: 5BK4) shows local DNA untwisting close to the interface of MCM2-7 complexes. (**b**) The MCM2-7 DH promotes local dsDNA destabilisation and melting *in silico*. An Ising model prediction for twist-induced dsDNA melting of the MCM2-7 DH (PDB ID: 5BK4) on origin dsDNA. A representative ARS sequence is shown above, and the DH-covered area is highlighted. (**c**-**e**) GC content analysis of (**c**) early, (**d**) late activating origins, and (**e**) randomised origin sequences. The 64 bp DH footprint (grey) and the 72 bp activating MCM2-7 DH footprint (orange bars) are highlighted, and their locations are indicated. (**f**) Schematic depiction of the 64 bp DH footprint and the experimentally observed 72 bp footprint observed here (see Fig. 5b). (**g**) Melting probability analysis of dsDNAs at early (blue) and late (green) origins, as well as random sequences (purple). A representative ARS sequence is shown above, the 64 bp DH footprint (grey) and the 72 bp activating MCM2-7 DH footprint (orange bars) are highlighted, and their locations are indicated.

Our ChIP-Exo analysis identified individual footprints for MCM2-7 DHs (Fig. 5a), and we observed that these DHs, in the context of the C-terminal Mcm2/5 linkage, remain bound close to their loading site during the G1-S transition (Fig. 5b and Supplementary Fig. 7)^19^. Therefore, we sought to characterise these DNA sequences *in silico* to identify properties that may be associated with helicase activation sites. The most prominent complex during helicase activation is the 72 bp footprint, which we interpret as a complex where MCM2-7 hexamers have been separated (Fig. 6c, Supplementary Fig. 1d). We calculated the GC content at early and late origins surrounding the DH-binding centre (Fig. 6d and f). Here, we observed a striking change in GC content across the MCM2-7 loading site, with AT maxima immediately upstream and downstream, and a GC maximum precisely at the centre (Fig. 6d and e). Interestingly, early and late origins exhibited similar properties in contrast to random yeast sequences (Fig. 6f). Interestingly, considering that both hexamers become separated during helicase activation, and we observed a 72 bp footprint in cells (Fig. 5c), we indicated the expected MCM2-7 hexamer position relative to each other and the GC content (see orange boxes in Fig. 6d, e, and g). Interestingly, the most AT-rich part appears to be located within the centre of each MCM hexamer. This could support DNA extrusion in S-phase, as AT-rich sequences melt more readily than GC-rich sequences, while the most GC-rich sequence is found between the two hexamers. This GC-rich maximum would also limit DNA unwinding in the context of the MCM2-7 DH, when both hexamers are fused together. As such, we now understand the GC-content across the yeast replication origins, which is characterised by a sharp “W” pattern right across the MCM2-7 loading site, which is similar between early and late origins (Fig. 6d and e). These sequence properties have the potential to regulate the extrusion of the lagging strand from the MCM2-7 complex during helicase activation.

### Initial DNA unwinding occurs near the MCM2-7 DH interface

Current models suggest that, during CDK-dependent helicase activation, 3 bp of DNA are initially destabilised within the central section of each MCM2-7 hexamer^22^ before Mcm10-dependent helicase activation induces complete strand separation within each MCM2-7 hexamer, resulting in >60bp of unwound DNA^24^. Consequently, it is thought that the two CMGs pass each other before processive DNA unwinding by the replication fork can start^10,24,25,44^. Although structural data have provided glimpses of this process, little is known about the location of initial DNA unwinding within the replication origin, whether prominent DNA unwinding intermediates exist, and how DNA unwinding occurs *in vivo* with a full complement of factors present, which are missing from reductionist cryo-EM structures. Understanding this process better, especially *in vivo*, is important for elucidating how replication origins function and are regulated, and to address whether different origins use diverse approaches for DNA unwinding or whether they all follow the same model.

Given that we observed Mcm10 recruitment by ChIP-MS and ChIP-qPCR (Fig. 4b, Supplementary Fig. 5f), we hypothesised that the Mcm2/5 linkage may arrest replisome assembly before or just after lagging-strand extrusion. Therefore, we aimed to directly detect DNA unwinding in cells. In bacteria and viruses, potassium permanganate footprinting has generated insights into the various stages of DNA unwinding at replication origins^45,46^. However, this analysis is absent for eukaryotes, especially across the genome. Therefore, we employed permanganate treatment and chromatin immunoprecipitation sequencing (PIP-seq) to identify MCM-associated initial DNA unwinding genome-wide (Fig. 7a)^47,48^. Cells harbouring the C-terminal Mcm2/5 linkage were arrested in G1-phase and released into S-phase in the presence of rapamycin (0-30 min). PIP-seq resulted in a robust increase in the detection of thymidine residues, as previously observed (Fig. 7b)^49^. By plotting the PIP-seq coverage at early origins, we detected increased hyperreactivity of thymidines during the G1-S transition (Fig. 7c). During G1-phase, when the MCM2-7 helicase is inactive, a centrally located baseline signal was detected in both strands (Fig. 7c). This signal is likely due to an MCM2-7-induced twist in the dsDNA (Fig. 6a and b), which disrupts nucleotide stacking and could facilitate permanganate attack. Interestingly, from the 15-minute time point onwards, ssDNA emerged specifically in the central section, i.e. at the MCM2-7 DH interface and from within the core origin DNA sequence (Fig. 7c). Observing thymidine hyperreactivity in the central section and its gradual expansion suggests that DNA extrusion from the CMG does occur in the context of the rapamycin-induced Mcm2/5 linkage. We suggest that Mcm10 is recruited and the lagging strand is extruded, but the linkage then interferes with the two CMGs passing each other, thereby restricting processive DNA unwinding until the linkage breaks.

**Fig. 7:**
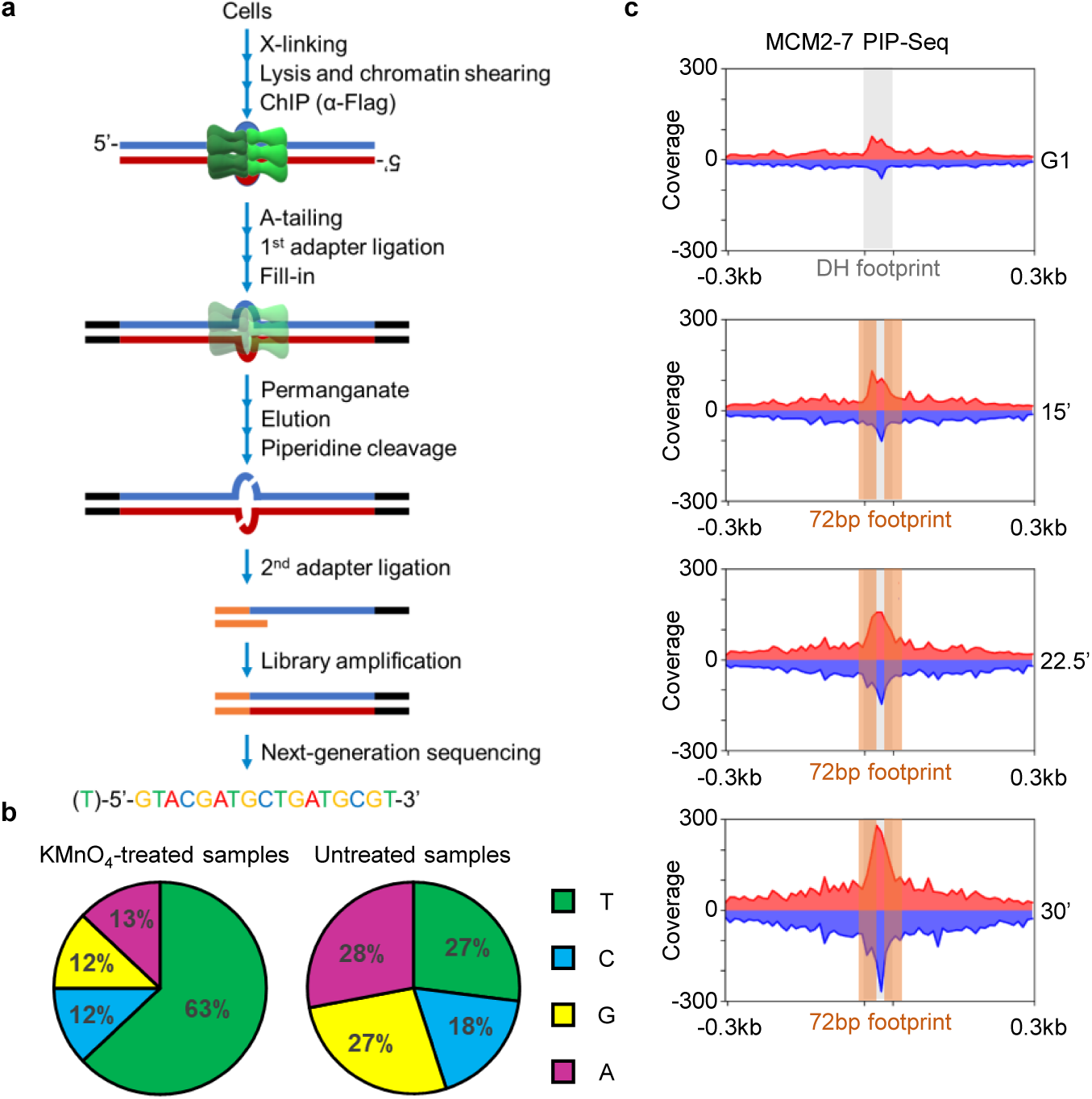
Initial unwinding is detected adjacent to the N-terminal interface of the MCM2-7 hexamers. ssDNA emerges from within the MCM2-7 DH. (**a**) Schematic overview of the PIP-Seq workflow. See Materials and Methods for details. (**b**). Permanganate-driven oxidisation of exposed thymidines defines the specificity of PIP-Seq. Distribution of permanganate-piperidine cleavage sites at different nucleotides. Potassium permanganate-treated samples (left) versus untreated samples (right). (**c**) PIP-Seq time course analysis of cells arrested in G1 and released into S-phase. Reactive thymidines from Mcm4 immunoprecipitated complexes were identified, and the coverage was plotted ±300 bp (forward strand in red and reverse strand in blue). The area covered by the DH was coloured grey, the area covered by the 72 bp split helicase activation complex was coloured orange, and signals were normalised to local GC content.

## DISCUSSION

This study provides detailed insight into helicase activation *in vivo*. Using a synthetic-biology approach, we linked the two hexamers of the MCM2-7 DH in a rapamycin-dependent manner to interfere with inter-hexamer rotation and double-hexamer separation. Consistent with a defect in helicase activation, we observed defects in cell proliferation and G1/S-phase cell cycle progression. Although CMG formation appeared normal, we observed delayed Sld3 release and delayed Mcm10 binding (Fig.1g and j). Recently, a cryo-EM study has shown that CDK-dependent helicase activation leads to CMG formation and inter-hexamer rotation (Supplementary Fig. 1d)^24^. Now, our data suggest that a latch-induced block in inter-hexamer rotation and/or separation (Fig. 1b) delays Mcm10 recruitment and interferes with Sld3 release and helicase activation (Fig. 8a-e). Thus, we provide the first evidence that inter-hexamer rotation/separation is required for the release of a rate-limiting helicase activation factor^50,51^. More detailed biochemical and cryo-EM analyses could pinpoint the specific replication factor, or combination of factors, required for Sld3 release.

**Fig. 8:**
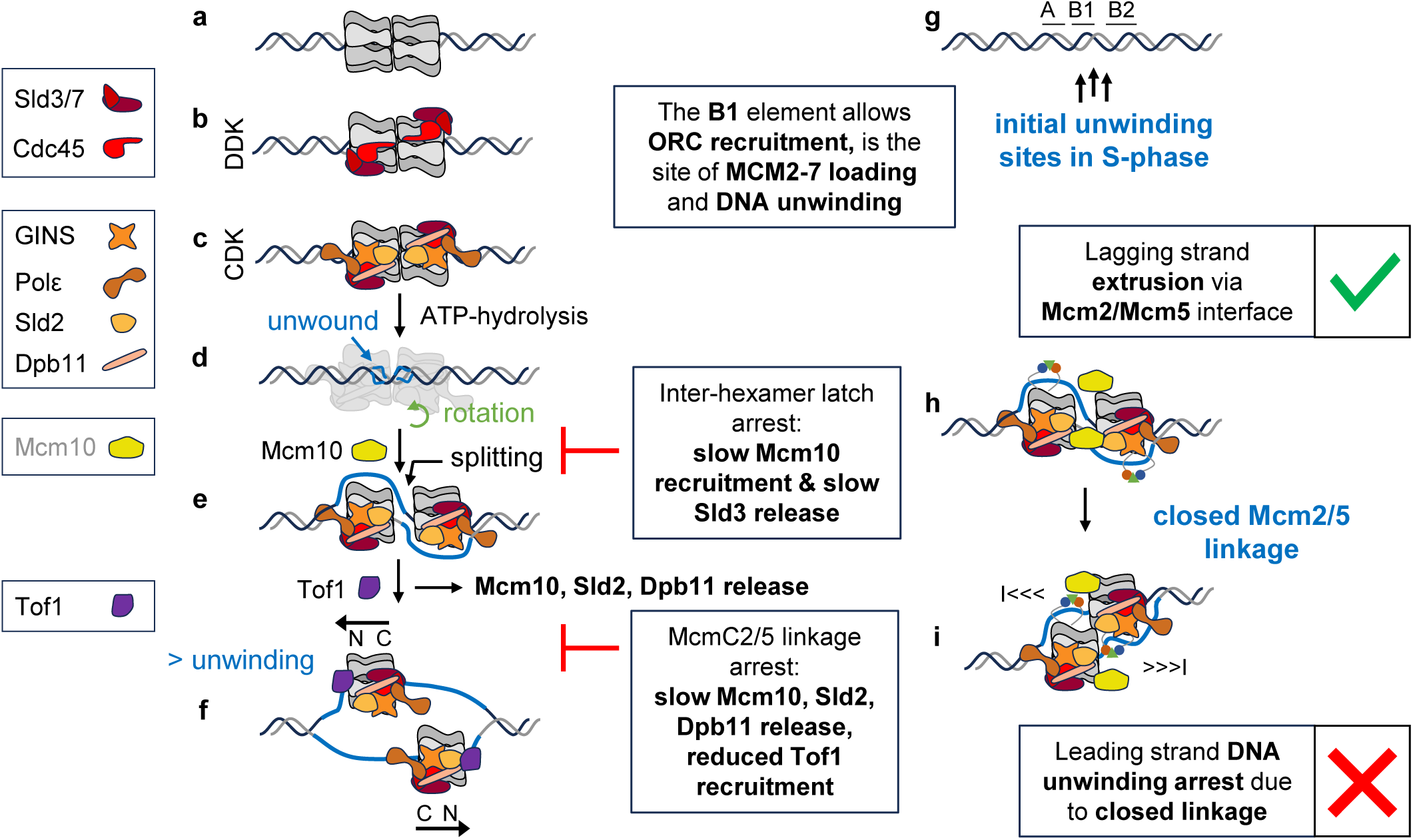
Model of DNA pathways and remodelling of the activating MCM2-7 DH. Model of MCM2-7 helicase activation in budding yeast. (**a**) MCM2-7 DH. (**b**) DDK activated MCM DH in complex with Sld3-Sld7. (**c**) CDK activity promotes the recruitment of helicase activation factors GINS, Polε, Sld2, and Dpb11. (**d**) Sld2-GINS-Dpb11-Polε-induced MCM2-7 ATPase-dependent activity results in the splitting of the DH and a rotation of the two CMGE complexes against each other. (**d**) Mcm10 recruitment promotes DNA extrusion and complete separation of the two hexamers. (**f**) Extended unwinding promotes the recruitment of Tof1, part of the replication fork protection complex. Loading factors that do not form part of the replication fork are shown in grey. **(g)** *In silico* prediction and *in vivo* PIP-seq analysis show initial DNA unwinding at the interface of the activating MCM2-7DH at their deposition site on origins. (**h-i**) Model of the McmC2/5 linkage arrest by the closed linkage formed by FRB (blue), FKBP (red), and rapamycin (green). (**h**) ssDNA is successfully extruded from the MCM2-7 hexamer, but confined by the Mcm2/5 linkage. (**i**) Progression of DNA unwinding is restricted, since the linkage of one hexamer hits the leading edge of the other hexamer, stopping the two hexamers from passing each other.

The MCM2-7 hexamer, as part of the CMG, must open to extrude the lagging strand during helicase activation (Fig. 8e and f). Whether one or all the intra-hexameric Mcm interfaces function in this process remains unknown. We found that the Mcm2/5 linkage, when incorporated at the N- or C-terminus, impedes replication fork assembly, but not ongoing DNA synthesis (Fig. 2 and 3, Supplementary Fig. 3 and 4). In the case of the Mcm3/5 interface, only the N-terminal linkage had an impact and not the C-terminal, which does not fit with the concept of ssDNA extrusion along the entire MCM complex. Notably, all other interface constructs failed to restrict DNA replication (Fig. 3a and b). When FRB, FKBP, and rapamycin link Mcm2 and Mcm5 at the C-terminus, we observed cell-cycle arrest, recruitment of most helicase-activation factors, including Mcm10, and initial DNA strand separation (Fig. 2, 3, 4, and 7). However, fork progression factors, such as Tof1 (Fig. 8f), were largely absent. We suggest that ssDNA extrusion is achieved, but that the linkage captures the extruded ssDNA strand (Fig. 8h). In turn, the linkage restrains rapid, progressive DNA unwinding, specifically preventing the two CMGs from passing each other (Fig. 8i). This is consistent with the appearance of ssDNA near the interface of the two MCM2-7 hexamers (Fig. 7c). Breakage of the linkage due to the helicase motor action, or the suboptimal ssDNA path confined by the linkage, may explain the slow progression of DNA synthesis. However, we note that the addition of rapamycin after helicase activation had no major impact on ongoing DNA synthesis (Supplementary Fig. 3b and c). Thus, we conclude that the Mcm2/5 interface represents the DNA exit gate during helicase activation.

Moreover, we uncover that the passing of the forks is required for complete assembly of the replication fork and, concomitantly, for the release of the helicase activation factors Sld2, Dpb11, and Mcm10, which were retained in the presence of rapamycin and the Mcm2/5 linkage (Fig. 4d, Supplementary Fig. 5). We note that this concept is consistent with the full replication fork speed being dependent on Tof1, with both forks passing each other and operating independently^52^. Now, our data place the successful release of the lagging strand from the CMG and the passage of the two forks as a core step in replication fork assembly, as blocking this process results in complexes resembling a late-replication fork intermediate that is missing key components of the replication fork, and helicase activation factors are still retained, although single-strand extrusion occurred.

Here, we have used PIP-seq to investigate the initial DNA unwinding process in cells (Fig. 7). Surprisingly, initial DNA destabilisation already occurs in G1-phase at the DH interface, likely due to MCM2-7-induced structural changes in the DNA^53^ (Fig. 6a and b). Then, during the G1-S transition, this destabilised region expands (Fig. 7c). Concomitantly, we observed an increased dumbbell-shaped footprint (Supplementary Fig. 7b), suggesting DH splitting. Thus, the data indicate that the linkage does not block DH splitting or initial DNA unwinding, but we suggest that the CMGs fail to pass one another, accumulating ssDNA between each CMG complex (Fig. 5c, 7c and 8h). Consistently, we observed a block in Tof1 recruitment (Fig. 4d) under the same conditions.

By aligning DNA sequences via their MCM2-7 footprint, we observed drastic changes in AT content and the melting probability of the underlying sequences (Fig. 6c-g). We suggest that the GC maximum at the centre of the DH footprint may regulate DNA unwinding during helicase activation (Fig. 6c-f). Consistent with this model, it has been shown that MCM2-7 DH displacement from its normal binding site impairs helicase activation^54,55^.

In the context of the CMG, it has been shown that any Mcm interface may open to bypass a DNA-protein crosslink (DPC)^28^. Crucially, the shape of the inner MCM channel prevents DNA from freely rotating within the complex; thus, the relative distance between the CMG and DPC determines at which MCM interface the DPC will land. However, in the context of helicase activation, the subunit interface opening depends on the relative positioning of the two hexamers once activation occurs. Because MCM2-7 DH formation establishes a fixed orientation between the two hexamers, we suggest that only the Mcm2/5 interface becomes strained when the two CMGs pass each other. Therefore, this model explains how helicase activation and CMG bypass of a DPC differ and may impose distinct requirements on the respective MCM interfaces involved in ring opening.

Interestingly, DNA unwinding initiated near the *ARS* B1 element of origins (Fig. 8g), the same site used for ORC binding and helicase loading^20,56^. Thus, every step of DNA replication initiation, from origin recognition to origin licensing and initial DNA unwinding, occurs in close proximity to the B1 element, representing a hub of the *S. cerevisiae* replication origin (Fig. 5b). Considering that ORC is binding across the B1-element, the ORC - B1-element interaction may serve as a placeholder for later events during initiation of DNA replication. It is possible that other factors, such as chromatin organisation, forkhead transcription factors and Sir2^57,58^, control access to the B1 element and, in this way, regulate origin function *in vivo*.

In summary, our data generate a comprehensive view of helicase activation *in vivo*. We identified the DNA footprint of the CMG during helicase activation, linked it to specific DNA sequences, and identified the origin sequences associated with initial DNA unwinding. The linkages indicate that inter-hexamer rotation and separation are coupled to Sld3 release, and highlight that the Mcm2/5 gate plays a distinct role during helicase activation, providing mechanistic insights into helicase activation and origin function from *in vivo* data.

## METHODS

### Yeast strains and plasmids

Yeast strains, plasmids, and DNA sequences used in this study are listed in the Supplementary Material. *Saccharomyces cerevisiae* strains (W303) were used throughout the manuscript and grown at 30 °C in full medium (YPD) if not stated otherwise. For designing the individual MCM2-7 linkages/latch construct, a rapamycin-resistant yeast strain (K11607) was used, where *FPR1* was depleted, and FRB/FKBP domains were integrated genomically in frame into the respective *MCM* locus in the highlighted position. All genomically tagged strains were integrated into their endogenous locus and confirmed by colony PCR and Western Blot, if applicable.

### Dot spots

To assay for growth phenotypes of different MCM2-7 linkage strains, cells were grown overnight and subsequently dot spotted in a 10-fold dilution series on plates containing full medium (YPD) with or without added rapamycin (100 nM or 1 µM, Santa Cruz). Plates were then incubated at 30° C for 2-4 days and analysed.

### Flow cytometry

Flow cytometry experiments, data recording, gating, and analyses were performed as described earlier^59^ with the following modifications: Typically, 0.4 mL of a culture (OD_600_ > 0.5, 4.5x 10^6^ cells/mL) was sufficient to yield 40,000+ gated events per well. If detection of newly synthesised DNA was desired, an appropriate concentration of 5-ethynyl-2′-deoxyuridine (EdU, 2 µM) was added to the medium upon S-phase release. Unless otherwise stated, all steps were performed at room temperature (RT). 0.4 mL of cells were fixed with 1 mL 100% ethanol in a 96-well deep well plate and kept at 4 °C sealed with a foil and stored 4 °C until used. Cells were washed twice with 500 µL 1xTBS (2,500 *g*, 2 min in 20 mM Tris, 150 mM NaCl, pH 7.6), the plate was decanted and hit gently onto a piece of paper to dry. Cells were resuspended in 500 µL 1xTBS supplemented with RNaseA (10 mg/mL, 1/400 dilution) and incubated at 50 °C for 1 h. Then, 4 µL Proteinase K was added and the plate was incubated at 50 °C for 1 h. Cells were spun for 2 min at 2,500 *g*, the cell pellet was resuspended in 500 μL of 1xTBS +3 % BSA and incubated for at least 15 min. For the click reaction, samples were spun (2,500 *g*, 2 min), the supernatant was discarded, and the cell pellet was resuspended in 40 μL of freshly prepared dye azide mix. Per well, 31.8 μL of ultrapure water, 4 μL 10x TBS, 2 μL of CuSO_4_ (100 mM), 0.2 μL of AlexaFluor 647 picolyl-azide (500 μM), and 2 μL of sodium ascorbate (200 mg/mL) were mixed. The click mix was distributed and the reaction incubated in the dark for 30-60 min. Cells were washed once with 500 μL of 1xTBS+ 3% BSA, then with 500 μL of 1xTBS. Finally, cells were resuspended in 150 μL (cycling cells) to 300 μL (remaining samples) of 1x TBS, and 100 µL of the cell suspension was transferred into a new U-shaped-bottom 96-well plate. Sample sonication separated mother and daughter cells (70 % output, 2 s on, 2 s off for 20 s for 8-tip horn). SYTOX™ Green (final [c]= 2 µM) was added to each well of the plate and mixed. Samples were protected from light and measured on a flow cytometer equipped to measure 96-well plates. Samples were measured on an Agilent Novocyte Quanteon with the following settings: flow rate 14 μL/min, sample volume: 50 µL, mixing at 1,500 rpm every 8 wells for one cycle. All data were analysed with FlowJo (v10.10.0, BD Bioscience), and an example of the gating strategy is provided (Supplementary Fig. 8).

### ChIP

ChIP-qPCR experiments were performed as described earlier^60^. In summary, *S. cerevisiae* cells were grown in 100 mL YPD from overnight cultures to OD_600_ 0.5 at 220 rpm and 30 °C followed by 2 hrs α-factor arrest (5 µg/mL) at 25 °C, 220 rpm. Cells were spun down (500 *g*, 2 min, RT) washed once with pre-warmed media and released in 100 mL warm media. For samples where FRB/FKBP linkage closure was desired media was supplemented with 100 nM or 1 µM rapamycin 0.5 hrs before releasing and upon release. Samples were withdrawn every 15 min and cross-linked with 1% formaldehyde (Sigma) for 15 min at RT with mild agitation. Cross-linking was stopped by the addition of 0.25 M glycine and incubation for 5 min at RT. Cells were harvested, washed with 1x TBS, and flash frozen in liquid nitrogen until use. Pellets were thawed on ice, resuspended in 0.8 mL of FA-Lysis buffer (low salt) (50 mM HEPES-KOH pH 7.5, 150 mM NaCl, 2 mM EDTA pH 8.0, 1% Triton X-100 (v/v), 0.1% Na-Deoxycholate (w/v) and 0.1% sodium dodecyl sulphate (w/v)), supplemented with a Protease inhibitor cocktail and mixed with equal amounts of glass beads. Cell lysis was done on an MP FastPrep-24 5G with 3x 45 sec cycles (7 m/s) and 2 min breaks on ice. Lysates were sonicated using a Bioruptor (Diagenode) for 3 times 15 minutes (30 sec on/off) with 5 min breaks on ice to yield chromatin size between 0.25 - 0.5 kbp. The lysate was cleared by centrifugation twice (5 min and 10 min, 13,000 rpm at 4 °C). After measuring and equalising protein amounts, samples were incubated with a specific antibody against the protein/tag of interest (anti-FLAG 1/300, Sigma, F-1804) for 1.5 hrs on a roller at 25 °C. Subsequently, 15 μL ProtG Dynabeads were added and the samples were incubated for 1 h at 25 °C; beads were collected on a magnetic rack and washed with buffer (2x FA-Lysis low salt, 2x FA-Lysis high salt [50 mM HEPES-KOH pH 7.5, 500 mM NaCl, 2 mM EDTA pH 8.0, 1% Triton X-100 (v/v), 0.1% sodium deoxycholate (w/v) and 0.1% sodium dodecyl sulphate (w/v)], 2x TLEND [10 mM Tris-HCl pH 8.0, 250 mM LiCl, 1 mM EDTA pH 8.0, 0.5% NP-40 (v/v) and 0.5% sodium deoxycholate (w/v)] and 1x TE. Next, DNA was eluted from beads using ChIP elution buffer (50 mM Tris-HCl pH 7.5, 10 mM EDTA pH 8.0 and 1% sodium dodecyl sulphate (w/v)) and vigorous shaking at 65 °C for 20 min. After the addition of Proteinase K, samples were incubated for 2 hrs at 37 °C, followed by 12-16 hrs of incubation at 65 °C. Samples were purified using the QIAquick PCR Purification kit (Qiagen) according to the manufacturer’s instructions and diluted 1/5 to 1/10 before qPCR on selected origins.

### ChIP-Exo 5.0

Cells were grown in 100 mL YPD from diluted overnight cultures to OD_600_ 0.6 at 220 rpm and 30 °C followed by two hours α-factor arrest (final conc.: 5 µg/mL) at 25 °C, 220 rpm. Cells were spun down (500 *g*, 2 min, RT), washed once with pre-warmed media and released in 100 mL warm media (25 °C), and time points were taken as indicated. Cells were crosslinked, washed, lysed, and sonicated as described for ChIP. After immunoprecipitation and washing as described above, samples on beads were resuspended in 10 mM Tris-HCl (pH 8.0), transferred into 0.2 mL PCR strips, and the remaining steps were performed using a ThermoMixer C (Eppendorf) with a 96-well block as follows (all reagents were purchased from NEB unless stated otherwise)^61^: A-tailing (each reaction in 50 μL; 15 U Klenow fragment (-exo), 5 μL NEB buffer 2, and 100 μM dATP) was performed for 30 min at 37 °C, 1050 rpm. Beads were collected on a magnetic stand, the supernatant discarded, and the beads washed with 150 μL cold 10 mM Tris-HCl (pH 8.0). First adaptor ligation (each reaction 45 μL; 1200 U T4 DNA ligase, 10 U T4 PNK, 4.5 μL 10x NEBNext Quick ligation reaction buffer, and 375 nM adaptor mix (ExA2_iNN / ExA2B)) was incubated for 1 h at 25 °C, 1050 rpm. Beads were collected on a magnetic stand, the supernatant discarded, and the beads washed with 150 μL cold 10 mM Tris-HCl (pH 8.0). Subsequently, a fill-in reaction was set up (each reaction 40 μL; 10 U phi29 DNA polymerase, 4 μL 10x phi29 reaction buffer, 0.2 mg/mL BSA, and 180 μM dNTPs) mixed with the beads and incubated for 20 min at 30 °C, 1050 rpm. Beads were collected on a magnetic stand, the supernatant discarded, and the beads washed with 150 μL cold 10 mM Tris-HCl (pH 8.0). After that, DNA on beads was digested using λ exonuclease (each reaction 50 μL; 10 U λ exonuclease, 5 μL 10x λ exonuclease reaction buffer, 5 μL Triton X-100 (1% v/v), and 2.5 μL DMSO (5% v/v)) for 30 min at 37 °C, 1050 rpm. Beads were collected on a magnetic stand, the supernatant discarded, and the beads washed with 150 μL cold 10 mM Tris-HCl (pH 8.0). DNA was eluted from beads (each reaction in 40 μL; 3 μL ProteinaseK ( > 10 mg/mL, Sigma), 25 mM Tris-HCl (pH7.5), 2 mM EDTA (pH 8.0), 200 mM NaCl, and 0.5% sodium dodecyl sulphate (w/v)) for 25 min, 65 °C, 1200 rpm, and decrosslinked overnight (15.5 hrs) in a PCR cycler at 65 °C. To prepare for the second adaptor ligation, DNA was cleaned up using AMPure magnetic beads (Beckman Coulter). Second adaptor ligation was set up in 20 μL reactions (1200 U T4 DNA ligase, 2 μL 10x T4 DNA ligase buffer, 375 nM adaptor mix (ExA1-58/ ExA1SSL-N5)) and incubated for 1 h at 25 °C, 1050 rpm before clean-up with AMPure beads. Finally, libraries were amplified in 25 μL reactions (2U Phusion Hot Start polymerase (Thermo Scientific), 5 μL 5x Phusion HF buffer, 200 μM dNTPs, and 500 nM of primers (P1.3 and P2.1)) mixed with 15 μL purified DNA. PCR conditions were as follows: 20 s, 98 °C initial denaturation and 18 cycles of 20 s, 98 °C, 60 s, 52 °C, and 60 s, 72 °C.

After amplification, DNA libraries were gel extracted (2% agarose gel, DNA with a molecular weight between 200-500 bp) using the NEB Monarch PCR and DNA Cleanup Kit according to the manufacturer’s instructions to allow elution with smaller volumes. Libraries were quantified using Qubit fluorometry (Thermo Fisher, dsDNA HS Assay) and qPCR (KAPA Illumina library quantification) before pooling and Illumina next-generation sequencing (HiSeq 2500). Bam/ bai files were visualised using the Integrative Genomics viewer^62^.

### ChIP mass spectrometry

Cells were grown in 600 mL YPD from overnight cultures to OD_600_ 0.6 at 220 rpm and 30 °C followed by 2 hrs α-factor arrest at 25 °C, 220 rpm. Cells were spun down (500 *g*, 2min, RT) washed once with pre-warmed media and released in 600 mL warm media. For samples where FRB/FKBP linkage closure was desired media was supplemented with 100 nM rapamycin 0.5 hrs before releasing and upon release. At the desired time-point post-release, samples were crosslinked and washed as described for ChIP. Each sample was resuspended in 1.6 mL 1x TBS, and split into two tubes, before being spun again (8,000 *g*, 4 min, 4 °C) and the pellets frozen in liquid nitrogen and stored at -80 °C. Pellets were thawed on ice and resuspended in 800 μL of FA lysis (low salt) buffer complemented with a protease inhibitor cocktail, and two pellets split between three tubes, with 800 µL of glass beads added to each. Cells were lysed using an MP Fast-Prep 24 5G for 3x 45 seconds at 7 m/s with 2 min breaks on ice. Lysates were combined into two tubes and volume adjusted to 1.2 mL each with 1.5x total protease inhibitor cocktail concentration. Sonication and centrifugation were done as described for ChIP. Protein content of the supernatant was measured, and total protein amounts equated between samples by dilution, with input samples taken for quality control. The extract was then incubated with 7.2 μL anti-FLAG antibody (Sigma, F-1804) for 1.5 hrs at RT with rotation. 26 μL protein G Dynabeads were added and samples incubated for 1 hr. Beads were collected on a magnetic rack and subjected to washing (2x FA-lysis (low salt), 2x FA-lysis (high salt), 2x TLEND and 1x TE buffer) before combining the tubes and elution with ChIP elution buffer as done for ChIP. Samples were de-crosslinked at 95 °C for 15 min and analysed by SDS-PAGE followed by silver stain.

Immunoprecipitations (IP) were processed by single-pot solid-phase-enhanced sample-preparation (SP3) technology^63^. Eluted samples were mixed with reconstitution buffer (50 mM HEPES pH 8.0, 1% sodium dodecyl sulphate (w/v) SDS, 1% Triton X-100 (v/v), 1% NP-40 (v/v), 1% Tween 20 (v/v), 1% sodium deoxycholate (w/v), 5 mM EDTA pH 8.0, 50 mM NaCl, 1% glycerol (v/v)). Proteins were reduced using 5 mM DTT, followed by alkylation using 20 mM chloroacetamide. Reduced and alkylated protein was transferred to a new tube containing 100% EtOH pre-mixed with Sera-Mag Speedbeads (GE Healthcare, 2 μL of each per sample). Reduced and alkylated protein was added to a final concentration of 60% EtOH. Protein-bound beads were rinsed three times with 80% EtOH using a magnetic rack, and after the final wash, beads were resuspended in digestion solution (100 mM ammonium bicarbonate pH 8.0) containing 1 ug of Trypsin Gold (Promega). After sonication to disperse the beads, digestions were incubated overnight (18 hrs) at 37 °C and dried in a centrifugal evaporator. Total protein digests were dissolved in 0.1% trifluoroacetic acid by shaking (1200rpm) for 30 min and sonication for 10 min, followed by centrifugation (14,000rpm, 5°C and 10 min). LC-MS/MS analysis was carried out in technical duplicates. Chromatographic separation was performed using an Ultimate 3000 RSLC nano liquid chromatography system (Thermo Scientific) coupled to an Orbitrap Q-Exactive mass spectrometer (Thermo Scientific) via an EASY-Spray source. For LC-MS/MS analysis peptide solutions were injected and loaded onto a trapping column (Acclaim PepMap 100 C18, 100μm × 2cm) for desalting and concentration at 8 μL/min in 2% acetonitrile, 0.1% trifluoroacetic acid. Peptides were then eluted on-line to an analytical column (EASY-Spray PepMap RSLC C18, 75μm × 75cm). Eluted peptides were analysed by the mass spectrometer operating in positive polarity using a data-dependent acquisition mode. Ions for fragmentation were determined from an initial MS1 survey scan at 70,000 resolution, followed by HCD (Higher-energy Collision Induced Dissociation) of the top 12 most abundant ions at a resolution of 17,500. MS1 and MS2 scan AGC targets were set to 3e6 and 5e4 for maximum injection times of 50 ms and 50 ms, respectively. A survey scan m/z range of 400 - 1800 was used, normalised collision energy set to 37% and charge exclusion enabled for unassigned and +1 ions. Dynamic exclusion was set to 30 seconds.

### Potassium permanganate ChIP-Sequencing (PIP-Seq)

PIP-Seq samples were prepared essentially as described earlier, with the following changes^47,48^. *S. cerevisiae* cells were grown in 150 mL YPD from overnight cultures to OD_600_ 0.6 at 220 rpm and 30 °C, followed by 2 hrs α-factor arrest at 25 °C, 220 rpm. Cells were spun down (500 *g*, 2min, RT) washed once with pre-warmed media and released in 150 mL warm media supplemented with 100 nM rapamycin, if FRB/FKBP linkage closure was desired. Cells were crosslinked, washed and lysed as described for ChIP. Lysates were sonicated using a Bioruptor (Diagenode) for 15 cycles 30 s on/off in a cooled water and cleared by centrifugation twice (5 min and 15 min, 13,000 rpm at 4°C). Samples were incubated with an anti-Flag antibody (1/300, Sigma, F-1804) for 1.5 hrs on a roller at 25 °C. Subsequently, 15 μL ProtG Dynabeads were added, and the samples were incubated for 1 h at 25 °C; beads were collected on a magnetic rack and washed with buffer (1x FA lysis low salt, 1x FA lysis high salt, 2x TLEND and 2x 10 mM Tris-HCL pH 8.0). A-tailing, first adapter ligation and Fill-in reaction were done as described for ChIP-Exo 5.0. After the Fill-in reaction, samples were washed with 10 mM Tris-HCl (pH 8.0) and resuspended in 90 μL of 10 mM Tris-HCl (pH 8.0) and 10 mM potassium permanganate. The reaction was placed on a shaker (1,050 rpm) for exactly 1 min at 25 °C before 90 μL of STOP solution (10 mM Tris-HCl pH 8.0, 40 mM EDTA pH 8.0, 0.7 M mercaptoethanol and 2% sodium dodecyl sulphate (w/v)) was added and treated DNA was eluted with 190 μL ChIP elution buffer and vigorous shaking at 65 °C for 20 min. After the addition of Proteinase K, samples were incubated for 2 hrs at 37 °C, followed by 12-16 hrs of incubation at 65 °C. DNA was cleaned up by phenol/chloroform extraction followed by ethanol precipitation. DNA pellets were resuspended in 90 μL TE buffer, 10 μL of piperidine was added, and the reaction was incubated at 90°C for 30 min. 300 μL ultrapure water was added, and the piperidine was extracted twice in a new tube with 800 μL isobutanol, a third time with 400 μL of isobutanol, and finally with 400 μL of chloroform. Following a standard ethanol precipitation of the treated DNA, second adapter ligation and DNA library amplification and quantification were done as described for ChIP-Exo 5.0, with the exception that the libraries were cleaned up using AMPure XP beads (Beckman Coulter) according to the manufacturer’s instructions. Libraries were quantified using Qubit fluorometry (Thermo Fisher, dsDNA HS assay) and qPCR (KAPA Illumina library quantification) before pooling and Illumina next generation sequencing. Bam/ bai files were visualised using the Integrative genomics viewer^62^.

### Bioinformatical Analysis

#### ChIP-Exo

The raw sequencing reads were assessed for quality using FastQC (ver. 0.11.5)^64^. ChIP-Exo specific quality control was performed using ChIPexoQual^65^. Alignment of reads was done with Bowtie2 aligner (ver. 2.3.5) using default settings^66^. Bigwig files were generated for visualising with a genome browser using deeptools (ver. 3.2.1)^67^.

Read distributions of ChIP-Exo data were visualised using CHASE using standard parameters^68^. Footprints were produced with ChExMix (ver. 0.45) using standard parameters and --kldivergencethres -3 or -10 and --nomotifs^69,70^.

#### PIP-Seq

Sequencing reads from PIP-Seq were quality assessed using FastQC (ver. 0.11.5) and aligned to sacCer3 genome using Bowtie2 aligner^64,66^. Separate tracks for each strand were generated using bedtools (ver. 2.27.1) and UCSC wigToBigWig tool^67,71^. Aligned data was also filtered to only include reads that have either T or C positioned immediately before the first base of the read. Aligned data was additionally shifted to visualise the base that is immediately before the first base of these reads. To account for non-reactive nucleotide bases, signals were normalised to the local GC content. All profile plots were generated using Deeptools (ver. 3.2.1)^72^.

#### GC content analysis

GC content plots were generated by visualising genome-wide GC content over early and late origins using Deeptools (ver. 3.2.1). Genome-wide GC content was obtained from UCSC in bigwig format^73^.

#### ChIP mass spectrometry

Data were processed using the MaxQuant software platform (ver. 1.6.2.3), with database searches carried out by Andromeda search engine against the Swissprot *Saccharomyces cerevisiae* database (ver. 20180503, number of entries: 6,721)^74^. A reverse decoy database approach was used at a 1% false discovery rate (FDR) for peptide spectrum matches. Search parameters were as follows: maximum missed cleavages set to 2, fixed modification of cysteine carbamidomethylation and variable modifications of methionine oxidation, protein N-terminal acetylation and asparagine deamidation. Label-free quantification (LFQ) was enabled with an LFQ minimum ratio count of 2. ‘Match between runs’ function was used with match and alignment time limits of 0.7 and 20 min respectively.

For the generation of volcano plots, replicates and relative protein abundances were analysed using the Perseus software platform (ver. 1.6.10.43)^75^. Proteins that were not present in at least three out of four replicates in at least one condition were excluded. Missing values for retained proteins were imputed from a normal distribution. Proteins associated with the Replication Initiation GO term were compared with the volcano plot function in Perseus, with an FDR of 0.1 and S0 of 0.1^76^.

For generating enrichment plots (Figure 4 and S6), mean LFQ values from IP samples were used to calculate enrichment of proteins in IP versus a shared set of negative control samples. Where proteins were present in the IP but not the negative controls, an imputed value equal to the mean of the lowest 10% LFQ values in the negative controls was used. Standard error (S.E.) in the enrichment was calculated by error propagation, and upper and lower bounds of one S.E. were plotted on the logarithmic scale as upper and lower error bars. An imputed value equal to the average intensity of the lowest 10% of proteins detected in the NC sample was used when proteins lacked intensity measurements in the negative control.

#### Ising model for the computation of denaturation profiles

Melting profiles for each of the 118 early and late origins, and the same number of random yeast sequences were computed based on an Ising model (available at https://github.com/XiPacha/Ising-model)^40^. In this model, every base pair is discretised to be either in a regular or in a denatured state, denoted by 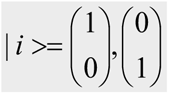. The Hamiltonian is determined by a bubble-initiation penalty (*ε* = 4.1*kcal* / *mol*), and sequence-specific base-pairing and base-stacking parameters (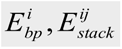, Tab 1)^77,78^:

**Table 1:** Base-stacking energies for all 10 dinucleotide steps. For base-pairing, we used *E_bp_* = 0.64/ 0.12 for GC/AT. All units are in kcal/mol.

|  |  |  |  |  |
| --- | --- | --- | --- | --- |
| AG | AC | AT | AA | TA |
| -1.44 | -2.19 | -1.72 | -1.49 | -0.57 |
| TC | TG | GG | GC | CG |
| -1.81 | -0.93 | -1.82 | -2.55 | -1.29 |

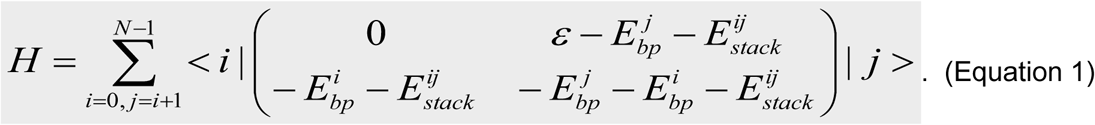

The partition sum is then obtained by transfer-matrix calculation:

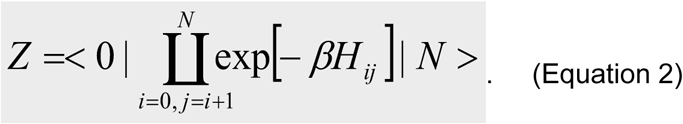

Melting probabilities were calculated through 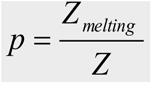, whereby 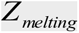 is derived from a Hamiltonian with requested base pairs constrained to the denatured state. Denaturation bubbles with a size up to 15 base pairs were allowed. By iterating over the entire sequence, complete melting profiles were obtained.

### Accounting for the DNA structure in complex with MCM2-7

The DNA structure in complex with the MCM2-7 double-hexamer (PDB: 5bk4) was parameterized with Curves+^53,79^. Consequently, the amount of unwinding for every turn (Δτ) of the DNA was quantified, and the associated deformation energy was computed by 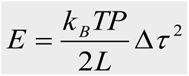. The contour-length was estimated to *L* = 10 · 0,34 nm, and the persistence length P to 100 nm. This energy term was added to the Hamiltonian (Equation 1), so that it promotes the denaturation initiation. On the other hand, energies associated with the unwinding of single-stranded regions were described by 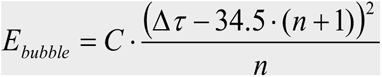. C denotes the force constant for unwinding of single-stranded regions, C = 0.79 cal/(mol deg²), and n the number of molten base-pairs. The remaining procedure was similar to the previous paragraph.

### Statistical Analysis

All ChIP-qPCR experiments presented are derived from at least three biological replicates. Values represent the average of the replicates, and the error bars are calculated as the standard deviation. ChIP-Exo and PIP-Seq experiments are derived from two biological replicates. ChIP-MS values are derived from four biological replicates and represent the mean with standard error derived error bars presented on a log(2) scale.

## SUPPLEMENTAL INFORMATION

Supplemental Information and Supplemental References can be found with this article online.

## DATA AVAILABILITY

All raw sequencing data that support the findings of this study have been deposited in the GEO repository with the accession codes GSE173525 and GSE173528. The mass spectrometry proteomics data have been deposited to the ProteomeXchange Consortium via the PRIDE partner repository with the dataset identifier PXD028809. Code for the Ising model computations is available at: https://github.com/XiPacha/Ising-model. Raw flow cytometry data annotated according to MIFlowCyt standards are available through FlowRepository^80,81^ (https://flowrepository.org) via the following experiment ID: FR-FCM-xxxx.

## ACKNOWLEDGEMENTS

We thank all members of the Speck lab for critical reading of the manuscript. We thank Anna Rogers and Conrad A. Nieduszynski for sharing unpublished plasmids pAR027, pAR047-049 and Karim Labib for sharing plasmid pKL255 used in this study. The authors would like to dedicate this manuscript to the late Dr. Holger Kramer, whose insights and mass-spectrometry support contributed to the development of this study. C.W. was supported by an EPSRC studentship (EP/N509486/1). This project received funding from the European Union’s Framework Programme for Research and Innovation Horizon Europe (2021-2027) under the Marie Sklodowska-Curie grant agreement no. 101109916 (to K.L.). M.Z. was funded by the German Research Foundation (DFG; grant ZA 153/28-1). H.D.U. was supported by the Deutsche Forschungsgemeinschaft (DFG, German Research Foundation), Project-ID: 393547839 - SFB 1361. C.S. was funded by the Biotechnology and Biological Sciences Research Council (BB/N000323/1 and BB/S001387/1), the Medical Research Council (MC_U120085811) and the Wellcome Trust (107903/Z/15/Z). L.M.R. was supported by the Deutsche Forschungsgemeinschaft (DFG, German Research Foundation), Project-ID: 505087959. We thank the IMB Media Lab for their support. In addition, we thank the IMB Flow Cytometry Core Facility, especially Stefanie Möckel and Stephanie Nick. Finally, we thank the LMS/NIHR Imperial Biomedical Research Centre Flow Cytometry Facility for support, and the MRC-LMS Genomics lab for Illumina sequencing.

## AUTHOR CONTRIBUTIONS

Conceptualization: C.S. and L.M.R.; Methodology: A.M., M.M.K., C.S., and L.M.R.; Software: A.M., K.L., and S.P.K; Validation: A.M., S.P.K., M.M.K, and L.M.R.; Formal analysis: C.W., K.L., S.P.K., and L.M.R.; Investigation: C.W., L.W., Au.M., A.M., V.R., K.L., and L.M.R.; Resources: C.W., L.W., Au. M., A.M., V.R., K.L., and L.M.R.; Data Curation: C.W., S.P.K., A.M., and M.M.K.; Writing: C.S. and L.M.R.; Visualization: C.W., K.L., S.P.K., and L.M.R.; Supervision: A.M., M.Z., M.M.K., H.D.U., C.S., and L.M.R.; Funding acquisition: A.M., M.Z., H.D.U., C.S., and L.M.R..

---

**Supplementary Fig. 1:**
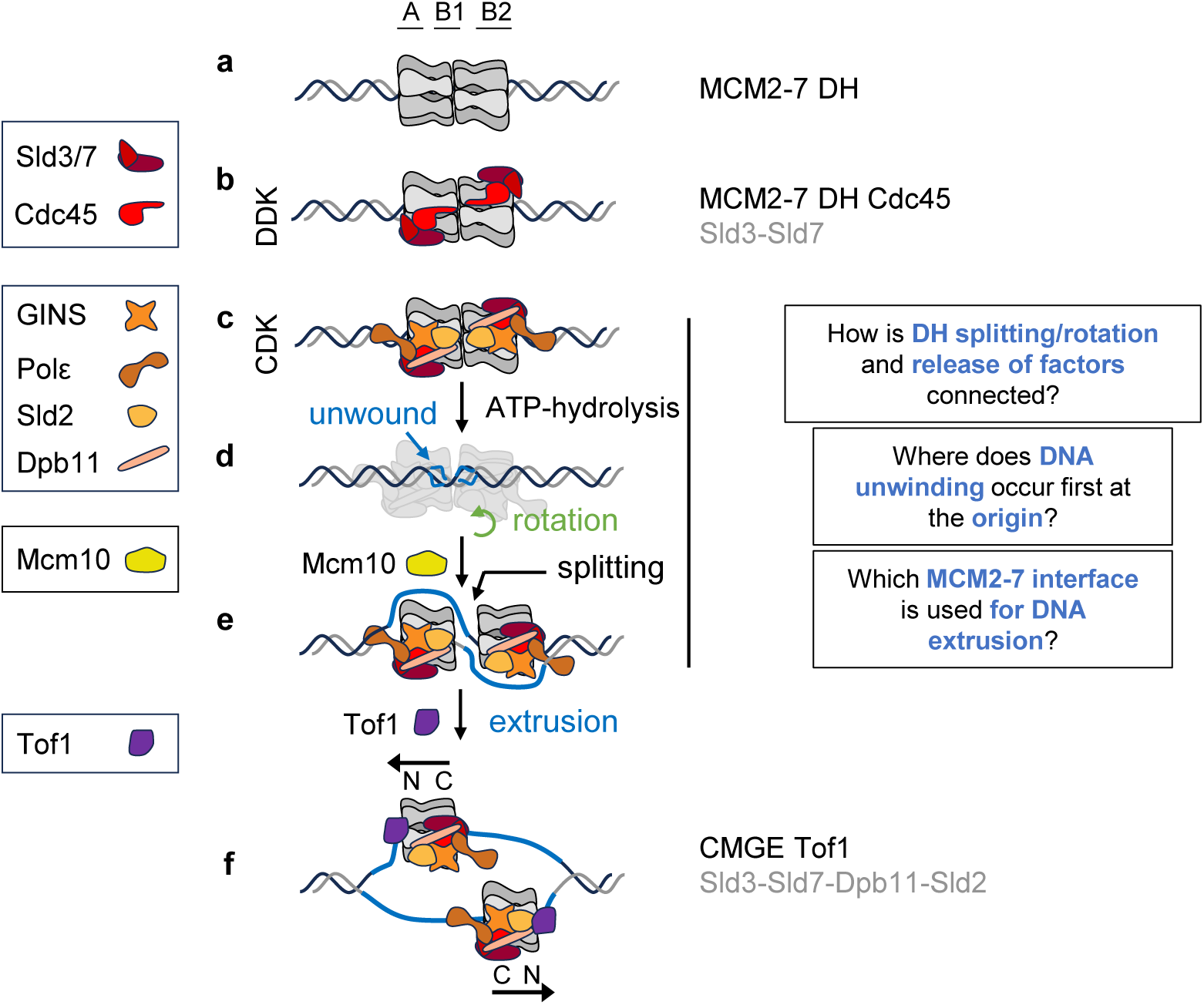
Helicase activation pathway. MCM2-7 helicase activation pathway in budding yeast. (**a**) MCM2-7 double-hexamer (DH). (**b**) DDK activated MCM DH in complex with Sld3-Sld7. (**c**) CDK activity promotes recruitment of helicase activation factors GINS, Polε, Sld2, and Dpb11. (**d**) Sld2-GINS-Dpb11-Polε-induced MCM2-7 ATPase-dependent activity results in the splitting of the DH and a rotation of the two CMGE complexes against each other. (**e**) Mcm10 recruitment promotes DNA extrusion and complete separation of the two hexamers. (**f**) Extended unwinding promotes the recruitment of Tof1, part of the replication fork protection complex. Loading factors that do not form part of the replication fork are shown in grey.

**Supplementary Fig. 2:**
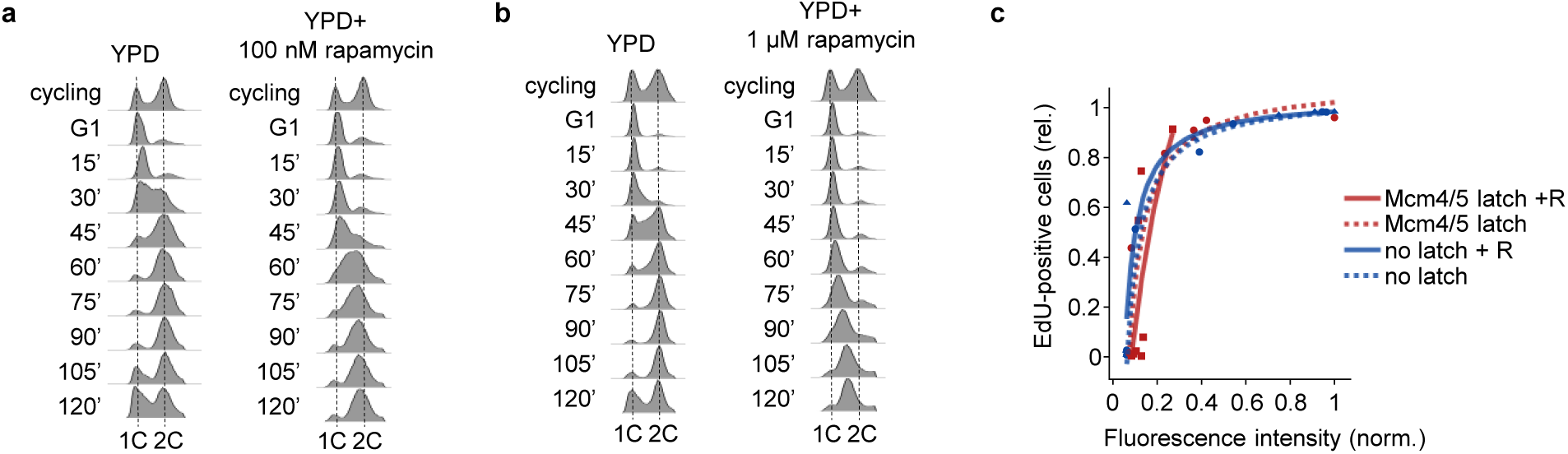
Analysis of the Mcm4/5 latch linkage. (**a**) Representative flow cytometry profiles of the inter-hexamer latch strain with and without rapamycin treatment. Cells were released into S-phase in the presence or absence of rapamycin after alpha-factor synchronisation. (**b**) As for (a), but with the addition of 1 µM rapamycin, showing improved rapamycin-dependent initiation delay. (**c**) Composite plot of an EdU incorporation time course of the Mcm4/5 inter-hexamer linkage in the (dotted lines) and presence of rapamycin (+R, 1 µM, solid lines). α–factor arrested cells were treated with rapamycin before release into S-phase for 90 minutes, and samples were taken every 15 min. Strains are colour-coded, and individual dots represent the relative percentage of EdU-positive cells and the geometric mean of EdU fluorescence intensity of at least 40,000 cells at a given time point (cyc, G1-90 min). Related to Fig. 1e and f.

**Supplementary Fig. 3:**
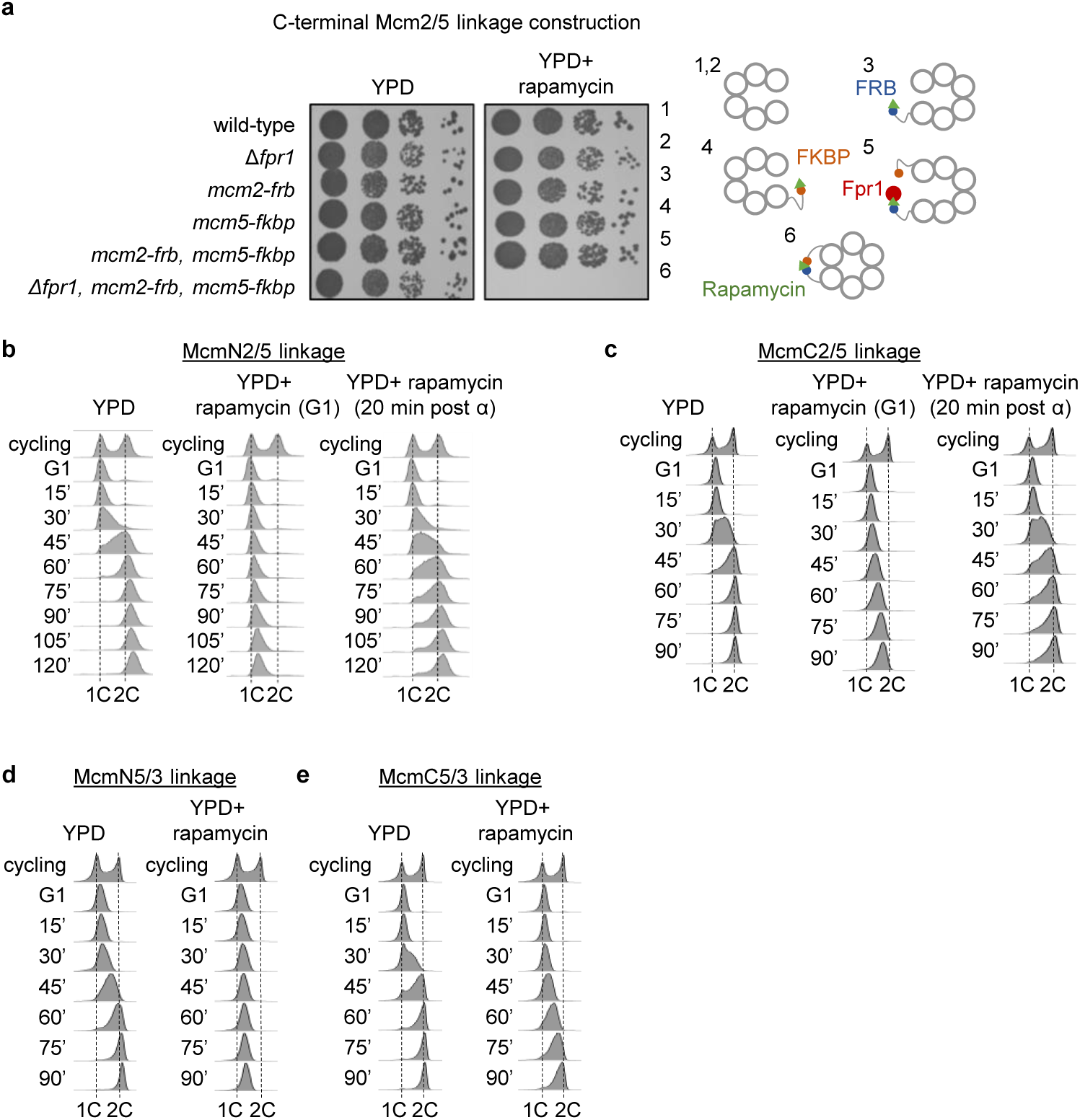
The Mcm2/5 linkage constructs result in specific cell cycle arrests. (**a**) The rapamycin-dependent lethal phenotype of the closed Mcm2/5 gate relies on a complete linkage construct. Dot spot assay of cells from various construction stages of the N-terminal Mcm2/5 linkage mutant on full media (YPD) or YPD-rapamycin (100 nM). Only the complete mutant strain harbouring all necessary genomic integrations is sensitive to rapamycin treatment. (**b** and **c**) The cell cycle arrest of the Mcm2/5 linkages is dependent on linkage closure before S-phase commitment. Representative flow cytometry profiles of a strain harbouring an (**b**) N-terminal Mcm2/5 linkage or (**c**) C-terminal Mcm2/5 linkage in the absence (left panel) and presence of rapamycin (100 nM, middle and right panels). Cycling cells were arrested for 2 hrs using α–factor and treated with rapamycin either during the G1-phase arrest (middle panel) or 20 minutes after release from the α–factor arrest (right panel). (**d** and **e**) Representative flow cytometry profiles of strains harbouring an (**d**) N-terminal or (**e**) C-terminal Mcm5/3 linkage in the absence (left panel) and presence of rapamycin (100 nM, right panel). Cycling cells were arrested for 2 hrs using α-factor and treated with rapamycin before release into S-phase.

**Supplementary Fig. 4:**
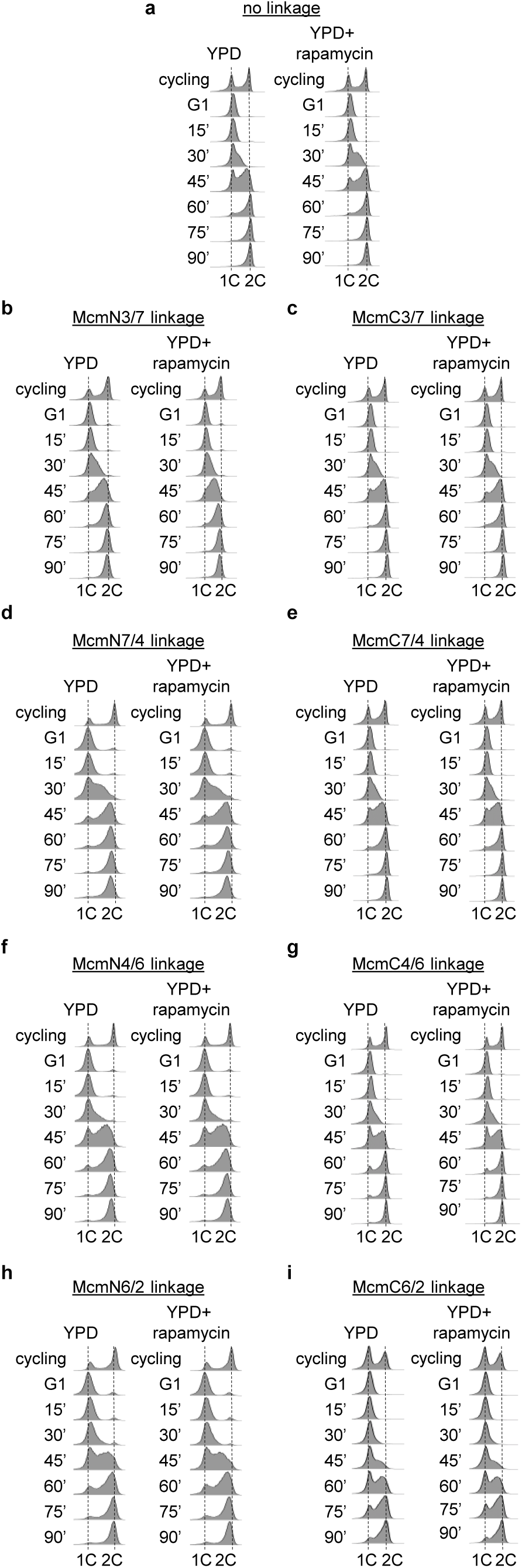
Flow cytometry profiles of the remaining N- and C-terminal Mcm2-7 linkages. Representative flow cytometry profiles of strains without an (**a**) FRB/FKBP linkage, (**b**, **d**, **f**, and **h**) N-terminal, or (**c**, **e**, **g**, and **i**) C-terminal Mcm linkages in the absence and presence of rapamycin (100 nM). Cycling cells were arrested for 2 hrs and treated with rapamycin before release into S-phase for 90 minutes. (**b** and **c**) Cytometry profiles for an (**b**) N-terminal or (**c**) C-terminal Mcm3/7 linkage strain. (**d** and **e**) Cytometry profiles for an (**d**) N-terminal or (**e**) C-terminal Mcm7/4 linkage strain. (**f** and **g**) Cytometry profiles for an (**f**) N-terminal or (**g**) C-terminal Mcm4/6 linkage strain. (**h** and **i**) Cytometry profiles for an (**h**) N-terminal or (**i**) C-terminal Mcm6/2 linkage strain.

**Supplementary Fig. 5:**
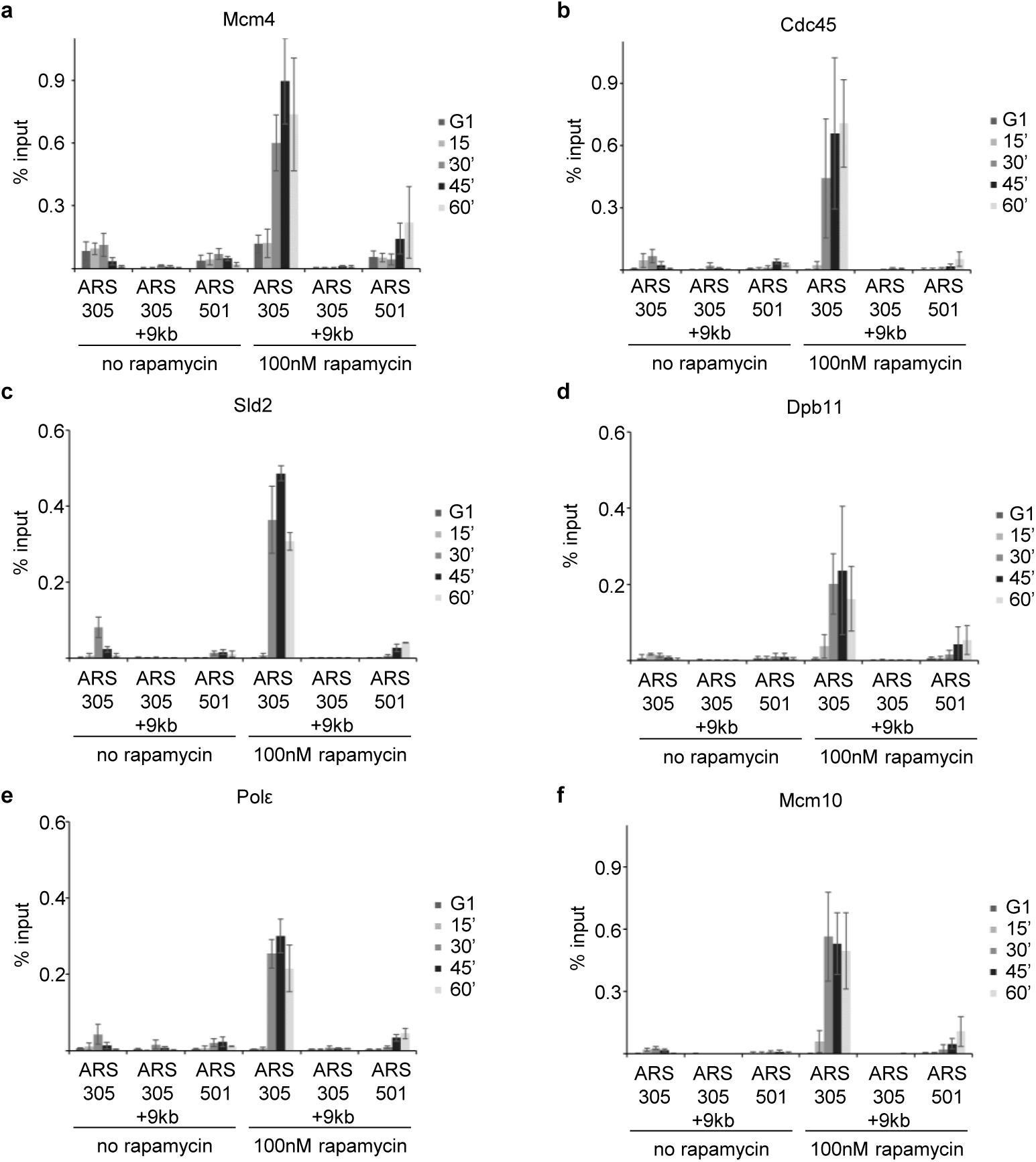
Recruitment of replication factors to origins in the presence of c-terminal Mcm2/5 linkage construct. ChIP-qPCR profiles of (**a**) Mcm4, (**b**) Cdc45, (**c**) Sld2, (**d**) Dpb11, (**e**) Polε, and (**f**) Mcm10 on *ARS305*, *ARS305*+9kb and *ARS501*. Tagged Mcm2/5 C-terminal linkage strains were treated with or without rapamycin (100nM) and harvested at the indicated timepoints after α-factor arrest and release at 25°C. Presented are the average and standard deviation of at least three biological replicates.

**Supplementary Fig. 6:**
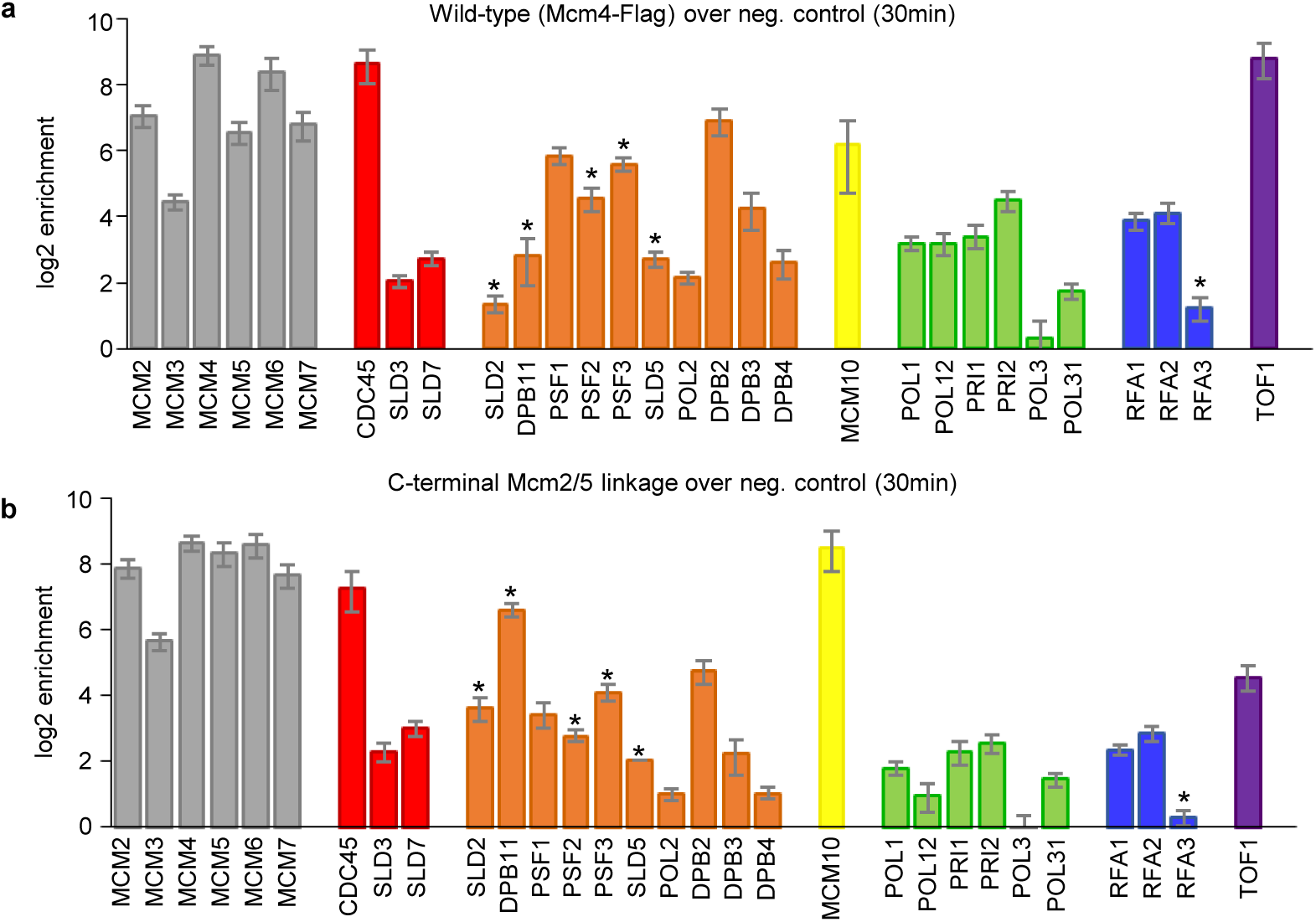
Protein composition of the WT and arrested McmC2/5 linkage complex. MCM2-7 complexes are enriched for proteins involved in replication activation and fork assembly. (**a**) Mcm4 ChIP-MS enrichment of core replisome factors in the wild-type strain, missing the linkage in G1-phase. Enrichment values are given as the log_2_ of the mean LFQ enrichment of subject proteins in Mcm4-5xFLAG strain (IP) over a negative control (NC – no Flag tag present) strain from four independent replicates. Asterisks mark proteins that lacked intensity measurements in the NC. Error bars represent the standard error (S.E.) of the non-logarithmic mean. (**b**) As in (**a**), showing Mcm4 ChIP-MS enrichment of core replisome factors in the Mcm2/5 C-terminal linkage strain released into S-phase for 30 min over a negative control (NC – no Flag tag present) strain. (**b**) is a copy of Fig. 4b, allowing for easier comparison of the data.

**Supplementary Fig. 7:**
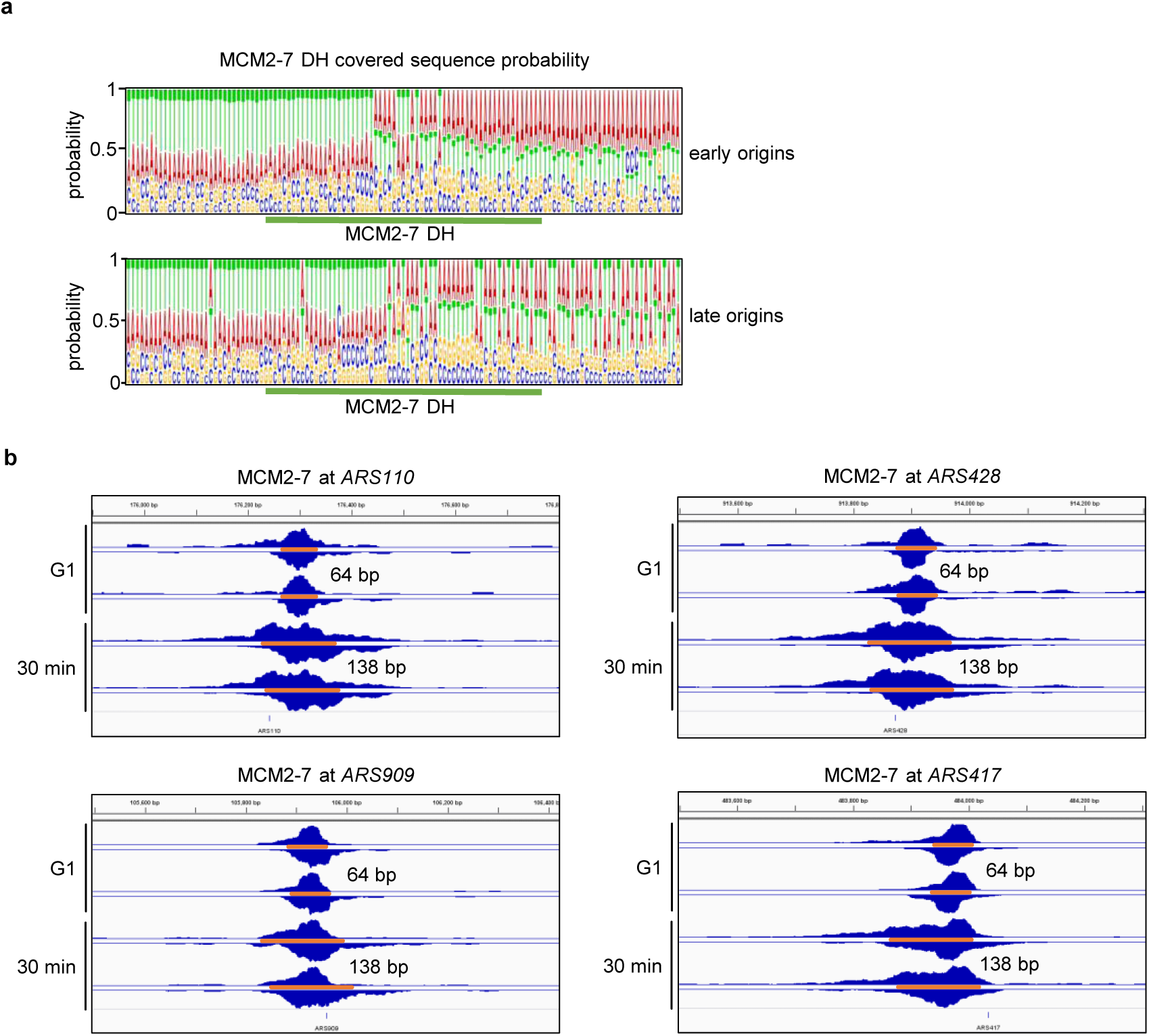
Genomic location of the closed McmC2/5 linkage construct. (**a**) Nucleotide frequency plots of MCM2-7 DH footprint centred at early (top panel) and late origins (lower panel) in the Mcm2/5 linkage strain. The area covered by MCM2-7 is highlighted (green). (**b**) ChIP-Exo 5.0 single traces of Mcm4 (G1 and 30min released, biol. duplicates) on origins (*ARS110*, *ARS428*, *ARS909*, and *ARS417*). Footprint sizes are indicated and highlighted with orange bars. Traces were visualised by IGViewer (ver. 2.4.14).

**Supplementary Fig. 8:**
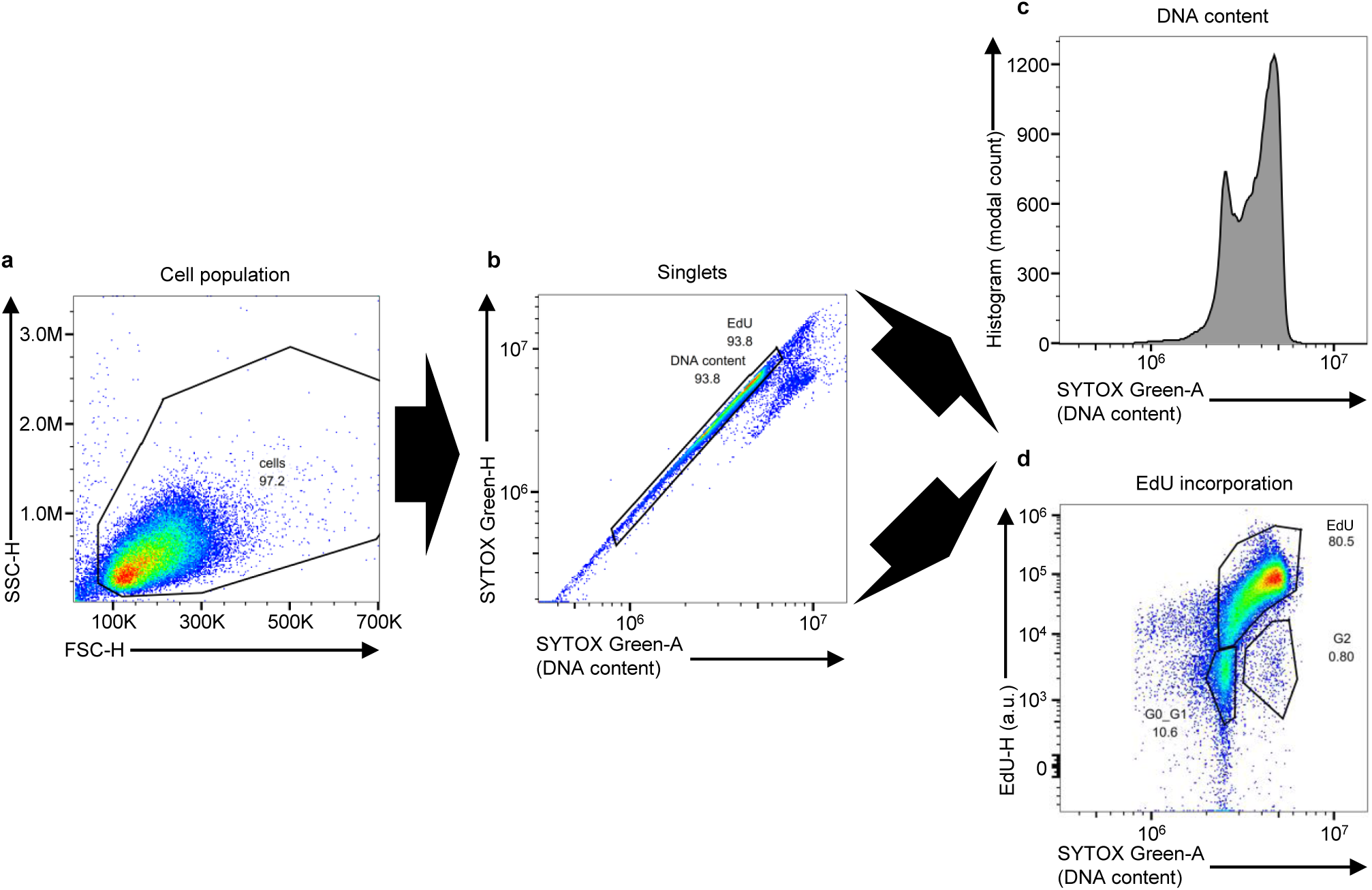
Gating strategy for DNA content and EdU incorporation analyses in budding yeast cells. (**a**) Cells of interest were identified via side scatter height (SSC-H) / forward scatter height (FSC-H) at linear scales (left panel). (**b**) Subsequently, doublets were excluded by SYTOX Green height (SYTOX Green-H) / SYTOX Green area (SYTOX Green-A) at linear scales (middle panel). (**c**) DNA content (SYTOX Green-A) was visualised by a one-dimensional histogram (488 nm laser and a 525/45 nm band pass (BP) filter) on a linear scale (top right panel). (**d**) EdU incorporation as a readout for *de novo* DNA synthesis was recorded using a 637 nm laser and a 667/30 nm BP filter (EdU height, relative fluorescence, and logarithmic scale) for AlexaFluor 647 versus SYTOX Green area (488 nm laser, 525/45 nm filter, and linear scale) (bottom right panel). Population gates are used to measure EdU incorporation at different cell cycle stages. The EdU gate includes both S-phase and G2-phase cells that have incorporated EdU. The numbers indicate the fraction of total analysed cells.

