## Supplementary material for "Mechanisms of MCM2-7 helicase activation and initial DNA melting at near base-pair resolution"

Content:

Supplementary Tables (1-3)

Supplementary References

### SUPPLEMENTARY TABLES

#### Supplementary table 1. Yeast strains

| Genotype | Identifier | Source |
|---|---|---|
| W303, *MATa, ade2-1, ura3-1, his3-11,15, trp1-1, leu2-3, 112, can1-100* | YC160 | Rothstein, R[1] |
| *MAT*a, *ade2-1, trp1-1, can1-100, leu2-3,112, his3-11,15, ura3, GAL, psi+, tor1-1* | K11607 | Nasmyth, K.[2] |
| *MATα, ade2-1 trp1-1 can1-100 leu2-3,112 his3-11,15, ura3 GAL, psi+, tor1-1, Δfpr1::NAT* | K14708 | Nasmyth, K.[2] |
| K11607*, ade2-1, trp1-1, can1-100, leu2-3,112, his3-11,15, ura3, GAL, psi+, tor1-1, Δfpr1::NAT* | yMR41 | This study |
| yMR41, *ChrVIII:202227-203265::pCCW12-hENT1_pTDH3-hsvTK, ChrXI:19786-20804::pTDH3-hsvTK, ChrXV:701610-702626::pTDH3-hsvTK* | yMR131 | This study |
| yMR41, *mcm2::mcm2-FRB-221-HisMX6* | YC446 | Samel et al., 2014[3] |
| K11607*, mcm5::mcm5-FKBP-72-kanMX6Δ* | YC450 | Samel et al., 2014[3] |
| K11607, *mcm2::mcm2-FRB-221-HisMX6, mcm5::mcm5-FKBP-72-kanMX6Δ* | YC451 | Samel et al., 2014[3] |
| yMR41, *mcm2::mcm2-FRB-221-HisMX6, mcm5::mcm5-FKBP-72-kanMX6Δ* | YC536 | This study |
| yMR41, *mcm2::mcm2-FRB-712-HisMX6, mcm5::mcm5-FKBP-591-kanMX6Δ* | YC525 | This study |
| yMR41, *mcm3::mcm3-FRB-37-HisMX6, mcm5::mcm5-FKBP-72-kanMX6* | YC778 | This study |
| yMR41, *mcm3::mcm3-FRB-645-HisMX6, mcm5::mcm5-FKBP-591-kanMX6* | YC796 | This study |
| yMR41, *mcm3::mcm3-FRB-37-HisMX6, mcm7::mcm7-FKBP-30-kanMX6Δ* | YC788 | This study |
| yMR131, *mcm3::mcm3-FRB-645-HisMX6, mcm7::mcm7-FKBP-630-kanMX6Δ* | yMR159 | This study |
| K14708, *mcm4::mcm4-FRB-252-HisMX6, mcm5::mcm5-FKBP-72-kanMX6Δ* | YC671 | This study |
| yMR41, *mcm4::mcm4-FRB-252-HisMX6, mcm7::mcm7-FKBP-30-kanMX6Δ* | YC771 | This study |
| yMR131, *mcm4::mcm4-FRB-739-HisMX6, mcm7::mcm7-FKBP-630-kanMX6Δ* | yMR158 | This study |

| | | |
|---|---|---|
| yMR41, *mcm4::mcm4-FRB-252-HisMX6, mcm6::mcm6-FKBP-124-kanMX6Δ* | YC772 | This study |
| yMR131, *mcm4::mcm4-FRB-739-HisMX6, mcm6::mcm6-FKBP-743-kanMX6Δ* | yMR153 | This study |
| yMR41, *mcm2::mcm2-FRB-221-HisMX6, mcm6::mcm6-FKBP-124-kanMX6Δ* | YC787 | This study |
| yMR131, *mcm2::mcm2-FRB-712-HisMX6, mcm6::mcm6-FKBP-743-kanMX6Δ* | yMR160 | This study |
| yMR131, *mcm2::mcm2-FRB-221-HisMX6, mcm5::mcm5-FKBP-72-kanMX6Δ* | yMR130 | This study |
| yMR131, *mcm2::mcm2-FRB-712-HisMX6, mcm5::mcm5-FKBP-591-kanMX6Δ* | yMR132 | This study |
| yMR131, *mcm3::mcm3-FRB-37-HisMX6, mcm5::mcm5-FKBP-72-kanMX6* | yMR134 | This study |
| yMR131, *mcm3::mcm3-FRB-645-HisMX6, mcm5::mcm5-FKBP-591-kanMX6* | yMR137 | This study |
| yMR131, *mcm3::mcm3-FRB-37-HisMX6, mcm7::mcm7-FKBP-30-kanMX6Δ* | yMR136 | This study |
| YC682, *mcm4::mcm4-FRB-252-HisMX6, mcm5::mcm5-FKBP-72-kanMX6Δ* | yMR150 | This study |
| yMR131, *mcm4::mcm4-FRB-252-HisMX6, mcm7::mcm7-FKBP-30-kanMX6Δ* | yMR138 | This study |
| yMR131, *mcm4::mcm4-FRB-252-HisMX6, mcm6::mcm6-FKBP-124-kanMX6Δ* | yMR133 | This study |
| yMR131, *mcm2::mcm2-FRB-221-HisMX6, mcm6::mcm6-FKBP-124-kanMX6Δ* | yMR135 | This study |
| YC536, *mcm4::MCM4-5xFlag::hphNT* | YC547 | This study |
| YC671, *mcm10::MCM10-5xFlag::hphNT* | YC675 | This study |
| YC671, *cdc45::CDC45-5xFlag::hphNT* | YC676 | This study |
| YC671, *sld3::SLD3-5xFlag::hphNT* | YC678 | This study |
| YC671, *pol2::POL2-5xFlag::hphNT* | YC679 | This study |
| YC671, *rfa1::RFA1-5xFlag::hphNT* | YC680 | This study |
| YC671, *mcm2::MCM2-5xFlag::hphNT* | YC682 | This study |
| YC525, *mcm10::MCM10-5xFlag::hphNT* | YC580 | This study |
| YC525, *cdc45::CDC45-5xFlag::hphNT* | YC584 | This study |
| YC525, *dpb11::DPB11-5xFlag::hphNT* | YC586 | This study |
| YC525, *mcm4::MCM4-5xFlag::hphNT* | YC588 | This study |
| YC525, *pol2::POL2-5xFlag::hphNT* | YC590 | This study |
| YC525, sld2*::SLD2-5xFlag::hphNT* | YC594 | This study |
| YC160, *mcm4::MCM4-5xFlag::hphNT* | YC779 | This study |

**Supplementary table 2. Plasmids**

| Details | Identifier | Source |
|---|---|---|
| pKL255 (5xFLAG-hphNT) | pCS762 | Labib, K. |
| pCas9 int-gRNA Ura3 | pAR027 | Nieduszynski, C. |
| pInt-ChrVIII hENT1-hsvTK | pAR047 | Nieduszynski, C. |
| pInt-ChrXI hsvTK | pAR048 | Nieduszynski, C. |
| pInt-ChrXV hsvTK | pAR049 | Nieduszynski, C. |
| pFA6a-*MCM2-FRB-221-HisMX6* | pCS642 | This study |
| pFA6a-*MCM2-FRB-712-HisMX6* | pCS756 | This study |
| pFA6a-*MCM3-FRB-37-HisMX6* | pCS1170 | This study |
| pFA6a-*MCM3-FRB-645-HisMX6* | pCS1185 | This study |
| pFA6a-*MCM4-FRB-252-HisMX6* | pCS1171 | This study |
| pFA6a-*MCM4-FRB-739-HisMX6* | pMR72 | This study |
| pFA6a-*MCM5-FKBP-72-kanMX6Δ* | pCS641 | This study |
| pFA6a-*MCM5-FKBP-591-kanMX6Δ* | pCS757 | This study |
| pFA6a-*MCM6-FKBP-124-kanMX6Δ* | pCS1172 | This study |
| pFA6a-*MCM6-FKBP-743-kanMX6Δ* | pMR73 | This study |
| pFA6a-*MCM7-FKBP-30-kanMX6Δ* | pCS1173 | This study |
| pFa6a-*MCM7-FKBP-630-kanMX6Δ* | pMR74 | This study |
| pRS316-*FPR1* | pMR68 | This study |

All FKBP and FRB proteins represent internal fusions into the Mcm proteins, with the number indicating the amino acid after which the fusion starts.

**Supplementary table 3. Oligos**

| Sequence 5’-3’ | Identifier | Source |
|---|---|---|
| **qPCR oligos** | | |
| AGCCTTCTTTGGAGCTCAAG | ARS305_fw | Reuter et al., 2024[4] |
| TTTGAGGAATTTCTTTTGAAGAG | ARS305_rev | Reuter et al., 2024[4] |
| AACTTGGCCTTATGTAGAATTTCTT | ARS305_+9fw | Reuter et al., 2024[4] |

| | | |
|---|---|---|
| AGCAATTCCACCGACCATAC | ARS305_+9rev | Reuter et al., 2024[4] |
| CTCATCATCATCCCCGGTA | ARS501_fw | Reuter et al., 2024[4] |
| CGTACACTAGCCCGTTGAGGT | ARS501_rev | Reuter et al., 2024[4] |
| **Genomic integration oligos** | | |
| GGCCTTTCACCTAAACTCGAGTATAAGCAAAAAATCAATCAAAACAAGTAATACGGATCCCCGGGTTAATTAA | Δfpr1_fw | This study |
| TACCATAAACATAAATAAAAAGCAGAAAGGCGGCTCAATTGATAGTACTTTGCTTGAATTCGAGCTCGTTTAAAC | Δfpr1_rev | This study |
| GTGAAGATCTTTCACCATTCCTGGAGAAGCTGACCTTGAGTGGATTGTTACGTACGCTGCAGGTCGAC | Cdc45-tag_fw | This study |
| ATATTCATATGCTGGTATATATGTACGACTAAATAATATAAATTTGATTAATCGATGAATTCGAGCTCG | Cdc45-tag_rev | This study |
| GGAAACTAAAGAAACTTCTGACGGTAGTGCCAGCGATCTTGAGATAATACGTACGCTGCAGGTCGAC | Mcm10-tag_fw | This study |
| CAAAAGTAATACCATTTTGGGGCCCTGAAAAGCACACCAATACTTATTTAATCGATGAATTCGAGCTCG | Mcm10-tag_rev | This study |
| GCTGAAGCCGACTATCTTGCCGATGAGTTATCCAAGGCTTTGTTAGCTCGTACGCTGCAGGTCGAC | Rfa1-tag_fw | This study |
| TTTCTCATATGTTACATAGATTAAATAGTACTTGATTATTTGATACATTAATCGATGAATTCGAGCTCG | Rfa1-tag_rev | This study |
| GTCCTTGGCGAGGGTGTAAGGAGATCAGTTCGCCTGAATAACCGTGTCCGTACGCTGCAGGTCGAC | Mcm4-tag_fw | Reuter et al., 2024[4] |
| TAATTAGTATTTATTAATTGTTACGCAGGGAATGATTGTAGTAGACAGCAATCGATGAATTCGAGCTCG | MCM4_delCintegr rev | Reuter et al., 2024[4] |
| GAAAAACCTATGAGACGACAGACAAGAAATCAGACAAAGGAATTAGATTCTCGTACGCTGCAGGTCGAC | Dpb11-tag_fw | This study |
| GATGGCGTATGTAAATGAATATCTTATAAAATTACGGACTACATTTCAATCGATGAATTCGAGCTCG | Dpb11-tag_rev | This study |
| GTATTACGGTTTTGATATATTATTGAGTTGTATTGCTGATTTGACCATACGTACGCTGCAGGTCGAC | Pol2-tag_fw | This study |
| TTTTCATGGTAAAGAGGCCATTGAACCTCGCGTTATATACTGCTTACTCAATCGATGAATTCGAGCTCG | Pol2-tag_rev | This study |
| CTAAAACTGCCAAAGAAAAACCGATTCTCTAATGGACGATGGGGAAGAAGGCGTACGCTGCAGGTCGAC | Sld2-tag_fw | This study |
| TAGGTTTCTATAAATTACAAATGTTTGTATTATTTACGCCATCACGCTCAATCGATGAATTCGAGCTCG | Sld2-tag_rev | This study |
| ATAGCTCAAAAAGGAGAGTAAGAAGACGTTTATTTGCTCCAGAATCCACACGTACGCTGCAGGTCGAC | Sld3-tag_fw | This study |
| GTTGGGTCAACTACTTGCCAGCGAAGGTGGTCTCGCAGTTTCTTATTCTAATCGATGAATTCGAGCTCG | Sld3-tag_rev | This study |
| CGCCGCCAACTCCGCAGGTCTTTCGCAATTTATACCTTGGGTCACCGTACGCTGCAGGTCGAC | Mcm2-tag_fw | This study |
| GAGAATTTTTTATCTTCATATCCAGATATTCGTAGGAATAACAAAGTTTTAATCGATGAATTCGAGCTCG | Mcm2-tag_rev | This study |

| | | |
|---|---|---|
| **Linkage construction oligos** | | |
| TGCTGCGGCAAATCCTAATG | MCM2aa647_fw | This study |
| ATGTCTGATAATAGAAGACGTAGAC | MCM2_ATG_fw | This study |
| CAGAGAATTTTTTATCTTCATATCCAGATATTCGTAGGAA TAACAAAGTTATCGATGAATTCGAGCTCG | MCM2_S2rev | This study |
| GAAAATGAAGGAGAAGATGATGAAGACCATGTCTTCGAA AAGTTCAACCCC | MCM3_aa645_fw | This study |
| ATGGAAGGCTCAACGGGATTTGATGG | MCM3_ATG_fw | This study |
| AAAAAAGCCAAGAATGGAAGTCTTTTAGTAAACATTCCTG TGACAATCGATGAATTCGAGCTCG | MCM3_S2rev | This study |
| GTCCTTGATAAGGTTGATGAGAAAAATGACAGAGAACTA GCCAAACAC | MCM4_aa739_fw | This study |
| ATGTCTCAACAGTCTAGCTCTCC | MCM4_ATG_fw | This study |
| TAATTAGTATTTATTAATTGTTACGCAGGGAATGATTGTA GTAGACAGCAATCGATGAATTCGAGCTCG | MCM4_S2rev | This study |
| GGATGACCATAATGAAGAACGTG | MCM5_aa591_fw | This study |
| ATGTCATTTGATAGACCGG | MCM5_ATG_fw | This study |
| CATGCAAACAAGTAGAAAAGGCGTCAAGCTAAGACTTTA TTGTTGATCGATGAATTCGAGCTCG | MCM5_S2rev | This study |
| TATTTTTTGTTATTCTTGATGACTGTAACGAAAAAATTGAT ACCG | MCM6aa743_fw | This study |
| ATGTCATCCCCTTTTCCAGCTGAC | MCM6_ATG_fw | This study |
| ACGATACTTGAAACGAAATATGAAAATCCGCAAGAGTGC ACTGAAATCGATGAATTCGAGCTCG | MCM6_S2rev | This study |
| GATATTCTCTTCTTAATGTTAGATATACCAAGTAGAGATG ACGACG | MCM7_aa630_fw | This study |
| ATGAGTGCGGCACTTCCATCAATTC | MCM7_ATG_fw | This study |
| AAAGAATAAAGAATGAAGGCCCTGTTGCTTTTTTTTTTAG AACTTATCGATGAATTCGAGCTCG | MCM7_S2rev | This study |
| CATCATTGGCACCAGCGTAATCAG | FRB_col_rev | This study |
| TTGCGTGAGGTGGTATAATACC | FKBP_col_rev | This study |
| **Cloning oligos** | | |
| ATTGCGGCCGCCACAGAAACTATTCAAC | FPR1_-600_NotI | This study |
| ATTGGATCCAGGCAGAAGGTAAAGG | FPR1_+500_BamHI | This study |

| ChIP-Exo oligonucleotides | | |
|---|---|---|
| 5-Phos CAA GCA GAA GAC GGC ATA CGA GAT TCGCCTTA GTG ACT GGA GTT CAG ACG TGT GCT CTT CCG ATC T | ExA2_iNN_701 | Reuter et al., 2024[4] |
| 5-Phos CAA GCA GAA GAC GGC ATA CGA GAT CTAGTACG GTG ACT GGA GTT CAG ACG TGT GCT CTT CCG ATC T | ExA2_iNN_702 | Reuter et al., 2024[4] |
| 5-Phos CAA GCA GAA GAC GGC ATA CGA GAT TTCTGCCT GTG ACT GGA GTT CAG ACG TGT GCT CTT CCG ATC T | ExA2_iNN_703 | Reuter et al., 2024[4] |
| 5-Phos CAA GCA GAA GAC GGC ATA CGA GAT GCTCAGGA GTG ACT GGA GTT CAG ACG TGT GCT CTT CCG ATC T | ExA2_iNN_704 | Reuter et al., 2024[4] |
| 5-Phos CAA GCA GAA GAC GGC ATA CGA GAT AGGAGTCC GTG ACT GGA GTT CAG ACG TGT GCT CTT CCG ATC T | ExA2_iNN_705 | Reuter et al., 2024[4] |
| 5-Phos CAA GCA GAA GAC GGC ATA CGA GAT CATGCCTA GTG ACT GGA GTT CAG ACG TGT GCT CTT CCG ATC T | ExA2_iNN_706 | Reuter et al., 2024[4] |
| 5-Phos CAA GCA GAA GAC GGC ATA CGA GAT GTAGAGAG GTG ACT GGA GTT CAG ACG TGT GCT CTT CCG ATC T | ExA2_iNN_707 | Reuter et al., 2024[4] |
| 5-Phos CAA GCA GAA GAC GGC ATA CGA GAT CCTCTCTG GTG ACT GGA GTT CAG ACG TGT GCT CTT CCG ATC T | ExA2_iNN_708 | Reuter et al., 2024[4] |
| 5-Phos CAA GCA GAA GAC GGC ATA CGA GAT AGCGTAGC GTG ACT GGA GTT CAG ACG TGT GCT CTT CCG ATC T | ExA2_iNN_709 | This study |
| 5-Phos CAA GCA GAA GAC GGC ATA CGA GAT CAGCCTCG GTG ACT GGA GTT CAG ACG TGT GCT CTT CCG ATC T | ExA2_iNN_710 | This study |
| 5-Phos CAA GCA GAA GAC GGC ATA CGA GAT TGCCTCTT GTG ACT GGA GTT CAG ACG TGT GCT CTT CCG ATC T | ExA2_iNN_711 | This study |
| 5-Phos CAA GCA GAA GAC GGC ATA CGA GAT TCCTCTAC GTG ACT GGA GTT CAG ACG TGT GCT CTT CCG ATC T | ExA2_iNN_712 | This study |
| 5-Phos CAA GCA GAA GAC GGC ATA CGA GAT TCATGAGC GTG ACT GGA GTT CAG ACG TGT GCT CTT CCG ATC T | ExA2_iNN_714 | This study |
| 5-Phos CAA GCA GAA GAC GGC ATA CGA GAT CCTGAGAT GTG ACT GGA GTT CAG ACG TGT GCT CTT CCG ATC T | ExA2_iNN_715 | This study |
| 5-Phos CAA GCA GAA GAC GGC ATA CGA GAT TAGCGAGT GTG ACT GGA GTT CAG ACG TGT GCT CTT CCG ATC T | ExA2_iNN_716 | This study |
| 5-Phos CAA GCA GAA GAC GGC ATA CGA GAT GTAGCTCC GTG ACT GGA GTT CAG ACG TGT GCT CTT CCG ATC T | ExA2_iNN_718 | This study |
| AAT GAT ACG GCG ACC ACC GAG ATC TAC ACT CTT TCC CTA CAC GAC GCT CTT CCG ATC T | ExA1-58 | Rossi et al., 2018[5] |
| GAT CGG AAG AGC ACA CGT CTG AAC TCC AGT CAC | ExA2B | Rossi et al., 2018[5] |
| NNN NNA GAT CGG AAG AGC G | ExA1-SSL_N5 | Rossi et al., 2018[5] |
| AAT GAT ACG GCG ACC ACC | P1.3 | Rossi et al., 2018[5] |

| | | |
|---|---|---|
| CAA GCA GAA GAC GGC ATA CGA G | P2.1 | Rossi et al., 2018[5] |
